# An N-terminal motif in NLR immune receptors is functionally conserved across distantly related plant species

**DOI:** 10.1101/693291

**Authors:** Hiroaki Adachi, Mauricio Contreras, Adeline Harant, Chih-hang Wu, Lida Derevnina, Toshiyuki Sakai, Cian Duggan, Eleonora Moratto, Tolga O Bozkurt, Abbas Maqbool, Joe Win, Sophien Kamoun

**Affiliations:** The Sainsbury Laboratory, University of East Anglia, Norwich Research Park, Norwich NR4 7UH, UK; Imperial College London, Department of Life Sciences, London, United Kingdom

## Abstract

The molecular codes underpinning the functions of plant NLR immune receptors are poorly understood. We used *in vitro* Mu transposition to generate a random truncation library and identify the minimal functional region of NLRs. We applied this method to NRC4—a helper NLR that functions with multiple sensor NLRs within a Solanaceae receptor network. This revealed that the NRC4 N-terminal 29 amino acids are sufficient to induce hypersensitive cell death. This region is defined by the consensus MADAxVSFxVxKLxxLLxxEx (MADA motif) that is conserved at the N-termini of NRC family proteins and ~20% of coiled-coil (CC)-type plant NLRs. The MADA motif matches the N-terminal α1 helix of Arabidopsis NLR protein ZAR1, which undergoes a conformational switch during resistosome activation. Immunoassays revealed that the MADA motif is functionally conserved across NLRs from distantly related plant species. NRC-dependent sensor NLRs lack MADA sequences indicating that this motif has degenerated in sensor NLRs over evolutionary time.

---

Plants have evolved intracellular immune receptors to detect host-translocated pathogen virulence proteins, known as effectors (Dodds and Rathjen, 2010; Jones et al., 2016; Kourelis and van der Hoorn, 2018). These receptors, encoded by disease resistance (*R*) genes, are primarily nucleotide-binding, leucine-rich repeat proteins (NLRs). NLR-triggered immunity (also known as effector-triggered immunity) includes the hypersensitive response (HR), a type of programmed cell death associated with disease resistance. NLRs are widespread across eukaryotes and have been described in animals and fungi in addition to plants (Jones et al., 2016). In contrast to other taxa, plants express very large and diverse repertoires of NLRs, with anywhere from about 50 to 1000 genes encoded per genome (Shao et al., 2016; Steuernagel et al., 2018). Genome-wide analyses have defined repertoires of NLRs (NLRome) across plant species (Shao et al., 2016). An emerging paradigm is that plant NLRs form receptor networks with varying degrees of complexity (Wu et al., 2018). NLRs have probably evolved from multifunctional singleton receptors—which combine pathogen detection (sensor activity) and immune signalling (helper or executor activity) into a single protein—to functionally specialized interconnected receptor pairs and networks (Adachi et al., 2019a). However, our knowledge of the functional connections and biochemical mechanisms underpinning plant NLR networks remains limited. In addition, although dozens of NLR proteins have been subject to functional studies since their discovery in the 1990s, this body of knowledge has not been interpreted through an evolutionary biology framework that combines molecular mechanisms with phylogenetics.

NLRs are multidomain proteins of the ancient group of Signal Transduction ATPases (STAND) proteins that share a nucleotide-binding (NB) domain. In addition to the NB and LRR domains, most plant NLRs have characteristic N-terminal domains that define three subgroups: coiled-coil (CC), CC_R_ or RPW8-like (RPW8) and toll and interleukin-1 receptor (TIR) (Shao et al., 2016). In metazoans, NLRs confer immunity to diverse pathogens through a wheel-like oligomerization process resulting in multiprotein platforms that recruit downstream elements, such as caspases (Qi et al., 2010; Zhou et al., 2015; Hu et al., 2015; Zhang et al., 2015; Tenthorey et al., 2017). In plants, NLRs also oligomerize through their N-terminal domains, notably the common and widespread CC domain (Bentham et al., 2018). The CC domain of some, but not all NLRs, triggers cell death when expressed as a truncated domain indicating that it can undertake oligomerization on its own (Maekawa et al, 2011; Casey et al, 2016; Cesari et al, 2016; Wróblewski et al., 2018). However, the molecular mechanisms that underpin CC-NLR activation and subsequent execution of HR cell death have remained largely unknown until very recently. In two remarkable papers, Wang et al. (2019a; 2019b) have significantly advanced our understanding of both the structural and biochemical basis of NLR activation in plants. They reconstituted the inactive and active complexes of the Arabidopsis CC-NLR ZAR1 (HOPZ-ACTIVATED RESISTANCE1) with its partner receptor-like cytoplasmic kinases (RLCKs) (Wang et al., 2019a; 2019b). Cryo-electron microscopy (cryo-EM) structures revealed that activated ZAR1 forms a resistosome—a wheel-like pentamer that undergoes a conformational switch to expose a funnel-shaped structure formed by the N-terminal α helices (α1) of the CC domains (Wang et al., 2019a; 2019b). They propose an engaging model in which the exposed α1 helices of the ZAR1 resistosome mediate cell death by translocating into the plasma membrane and perturbing membrane integrity similar to pore-forming toxins (Wang et al., 2019b). However, whether the ZAR1 model extends to other CC-NLRs is unknown. One important unanswered question is the extent to which the α1 helix “death switch” occurs in other CC-NLRs (Adachi et al., 2019b).

Although ZAR1 is classified as a singleton NLR that detects pathogen effectors without associating with other NLRs, many plant NLRs are interconnected in NLR pairs or networks (Wu et al., 2018; Adachi et al., 2019a). Paired and networked NLRs consist of sensor NLRs that detect pathogen effectors and helper NLRs that translate this effector recognition into HR cell death and immunity. In the Solanaceae, a major phylogenetic clade of CC-NLRs forms a complex immunoreceptor network in which multiple helper NLRs, known as NLR-REQUIRED FOR CELL DEATH (NRC), are required by a large number of sensor NLRs, encoded by *R* gene loci, to confer resistance against diverse pathogens, such as viruses, bacteria, oomycetes, nematodes and insects (Wu et al., 2017). These proteins form the NRC superclade, a well-supported phylogenetic cluster divided into the NRC helper clade (NRC-helpers or NRC-H) and a larger clade that includes all known NRC-dependent sensor NLRs (NRC-sensors or NRC-S) (Wu et al., 2017). The NRC superclade has expanded over 100 million years ago (Mya) from an NLR pair that diversified to up to one-half of the NLRs of asterid plants (Wu et al., 2017). How this diversification has impacted the biochemical activities of the NRC-S compared to their NRC-H mates is poorly understood. For example, it’s unclear how the ZAR1 conceptual framework applies to more complex NLR configurations such as the NRC network (Adachi et al., 2019b).

This paper originates from use of the *in vitro* Mu transposition system to generate a random truncation library and identify the minimal region required for CC-NLR-mediated cell death. We applied this method to NRC4—a CC-NLR helper of the NRC family that is genetically required by a multitude of NRC-S, such as the potato late blight resistance protein Rpi-blb2, to cause HR cell death and confer disease resistance (Wu et al., 2017). This screen revealed that the N-terminal 29 amino acids of NRC4 are sufficient to induce cell death. Remarkably, this region is about 50% identical to the N-terminal ZAR1 α1 helix, which undergoes the conformational “death switch” associated with the activation of the ZAR1 resistosome (Wang et al., 2019b). Computational analyses revealed that this region is defined by a motif, following the consensus MADAxVSFxVxKLxxLLxxEx, which we coined the “MADA motif”. This sequence is conserved not only in NRC4 and ZAR1 but also in ~20% of all CC-NLRs of dicot and monocot species. Motif swapping experiments revealed that the MADA motif is functionally conserved between NRC4 and ZAR1, as well as between NLRs from distantly related plant species. Interestingly, NRC-S lack N-terminal MADA sequences, which may have become non-functional over evolutionary time. We conclude that the evolutionarily constrained MADA motif is critical for the cell death inducing activity of CC domains from a significant fraction of plant NLR proteins, and that the “death switch” mechanism defined for the ZAR1 resistosome is probably widely conserved across singleton and helper CC-NLRs.

## Results

### Mu mutagenesis of NRC4 reveals a short 29 amino acid N-terminal region that is sufficient for induction of HR cell death

The N-terminal CC domain of a subset of CC-NLR proteins can mediate self-association and trigger HR cell death when expressed on its own (Bentham et al., 2018). However, to date truncation experiments have been conducted based on educated guesses of domain boundaries (Maekawa et al., 2011; Casey et al., 2016; Cesari et al., 2016; Wróblewski et al., 2018). Moreover, one amino acid difference in the length of the assayed truncation can affect cell death inducing activity (Casey et al., 2016). Therefore, we designed an unbiased truncation approach using bacteriophage Mu *in vitro* transposition system to randomly generate a C-terminal deletion library of the helper NLR NRC4. By using a custom-designed artificial transposon (Mu-STOP transposon) that carries staggered translation stop signals at Mu R-end (Poussu et al., 2005), we targeted the full-length coding sequence of the NRC4 autoactive mutant, NRC4^D478V^, (referred to from here on as NRC4^DV^). We generated a total of 65 truncated NRC4^DV^::Mu-STOP variants and expressed these mutants in *Nicotiana benthamiana* leaves using agroinfiltration (Figure 1A). Remarkably, only a single truncate carrying the N-terminal 29 amino acids triggered visible cell death in *N. benthamiana* leaves (Figure 1B; Supplementary file 1). To validate this phenotype, we expressed NRC4 N-terminal 29 amino acids (NRC4_1-29_) fused with the yellow fluorescent protein (YFP) at the C-terminus in *N. benthamiana* leaves (Figure 2A). NRC4_1-29_-YFP triggered a visible cell death response, although the cell death intensity was weaker than that of the full-length NRC4^DV^-YFP (Figure 2B and 2D).

**Figure 1.**
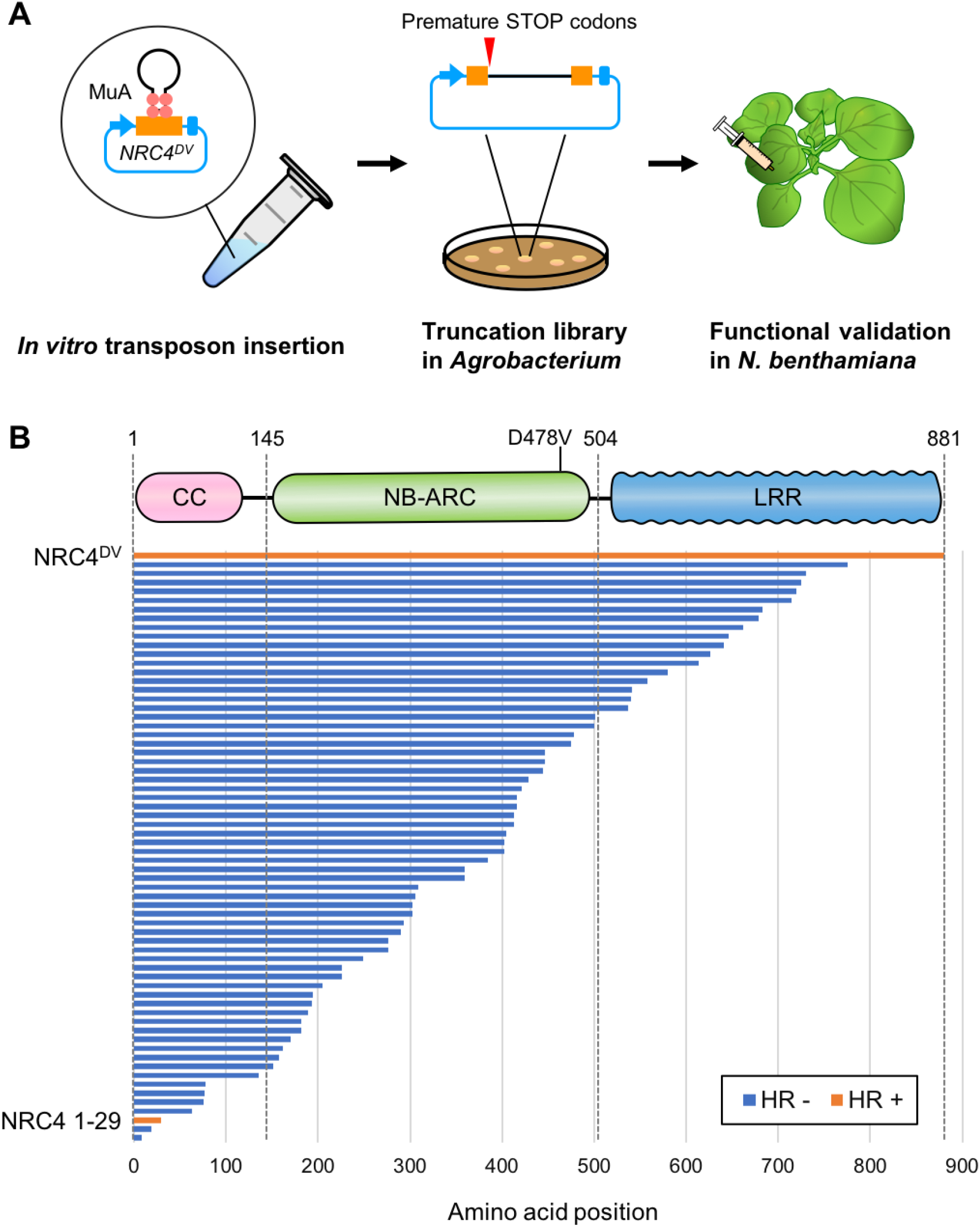
Transposon-based truncation mutagenesis reveals a short 29 amino-acid region sufficient for NRC4-mediated cell death. (**A**) Overview of the strategy for transposon-based C-terminal random truncation of NRC4 proteins. Hairpin Mu-STOP transposon and MuA proteins forming Mu transpososome were used for *in vitro* transposition into target plasmid. The truncation libraries (NRC4^DV^::Mu-STOP) were transformed into *Agrobacterium* for transient expression in *N. benthamiana* leaves. The tube, petri dish and syringe are not drawn to scale. (**B**) NRC4_1-29_::Mu-STOP triggers cell death in *N. benthamiana* leaves. In total, 65 truncated variants of NRC4^DV^ were expressed in *N. benthamiana* leaves, and the cell death activities are described as cell death induction (orange, HR+) and no visible response (blue, HR-).

**Figure 2.**
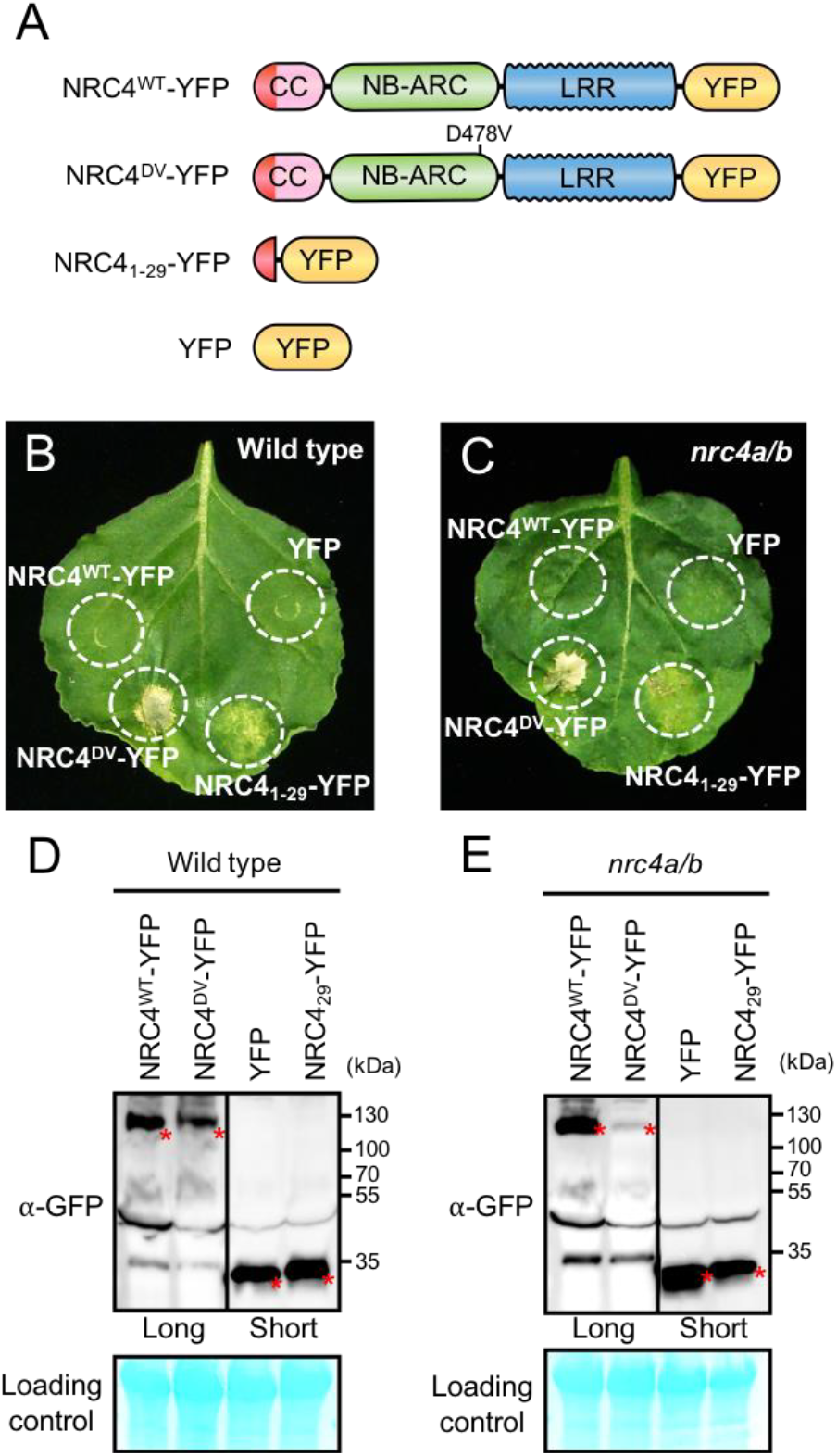
NRC4_1-29_-YFP induces cell death in *Nicotiana benthamiana* independently of endogenous NRC4. (**A**) Schematic representation of wild-type NRC4-YFP (NRC4^WT^-YFP) and the variants used for the *in planta* expression assays. The colour code is: red represents NRC4 1 to 29 amino acid region. (**B**) NRC4_1-29_-YFP triggers cell death in wild-type *N.* benthamiana leaves. NRC4^WT^-YFP, NRC4^DV^-YFP, NRC4_1-29_-YFP and YFP were co-expressed with the gene silencing suppressor p19 and photographed at 7 days after agroinfiltration. (**C**) NRC4_1-29_-YFP triggers cell death in *N. benthamiana* independently of endogenous NRC4. Leaves of *N. benthamiana nrc4a/b* line were used for agroinfiltration assays as described in B. (**D** and **E**) *In planta* accumulation of NRC proteins. Anti-GFP immunoblots of NRC4^WT^-YFP, NRC4^DV^-YFP, NRC4_1-29_-YFP and YFP expressed in *N. benthamiana* wild-type and *nrc4a/b* mutant. Total proteins were prepared from wild-type and *nrc4a/b N. benthamiana* leaves at 1 day after agroinfiltration. Given that the full-length NLRs accumulate at much lower levels than the shorter peptide, we showed different exposures as indicated by the black line. Red asterisks indicate expected band sizes.

To determine whether NRC4_1-29_-YFP requires the endogenous *N. benthamiana* NRC4 to trigger cell death, we expressed this fusion protein in mutant *nrc4a/b* plants that carry CRISPR/Cas9-induced mutations in the two NRC4 genes *NRC4a* and *NRC4b* (Figure Supplement 1; see methods). In these plants, NRC4_1-29_-YFP still induced cell death indicating that the activity of the N-terminal 29 amino acids of NRC4 is independent of a full-length NRC4 protein (Figure 2C and 2E).

### NRC4 carries N-terminal sequences that are conserved across distantly related CC-NLRs

Our finding that the N-terminal 29 amino acids of NRC4 are sufficient to trigger cell death prompted us to investigate the occurrence of this sequence across the plant NLRome. We first compiled a sequence database containing 988 putative CC-NLRs and CC_R_-NLRs (referred to from here on as CC-NLR database, Supplementary file 2) from 6 representative plant species (Arabidopsis, sugar beet, tomato, *N. benthamiana*, rice and barley) amended with 23 functionally characterized NLRs. Next, we extracted their sequences prior to the NB-ARC domain (Supplementary file 3). These sequences were too diverse and aligned poorly to each other to enable global phylogenetic analyses. Therefore, to classify the extracted N-terminal sequences based on sequence similarity, we clustered them into protein families using Markov cluster (MCL) algorithm Tribe-MCL (Enright et al., 2002) (Figure 3A). The 988 proteins clustered into 59 families of at least two sequences (tribes) and 43 singletons (Supplementary Table 1). The largest tribe, Tribe 1, consists of 219 monocot NLRs, including MLA10, Sr33, Sr50, the paired Pik and Pia (RGA4 and RGA5) NLRs, and 7 dicot NLRs notably RPM1 (Figure 3B). Tribe 2, the second largest tribe with 102 proteins, consists primarily of dicot proteins (93 out of 102) but still includes 9 monocot NLRs. Interestingly, Tribe 2 grouped NRC-H proteins, including NRC4, with well-known CC-NLRs, such as ZAR1, RPP13, R2 and Rpi-vnt1 indicating that these proteins share similarities in their CC domains (Figure 3B).

**Figure 3.**
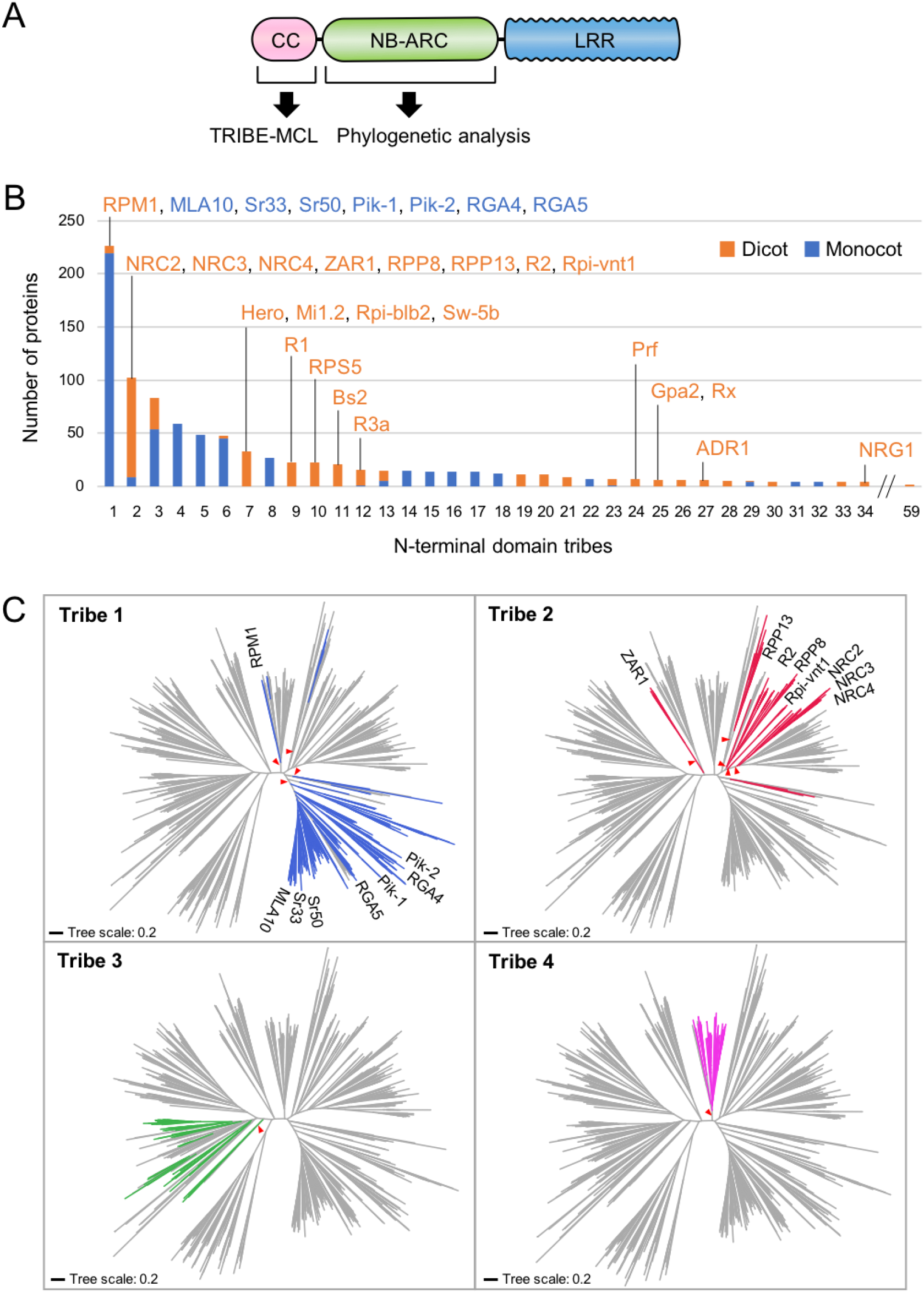
NRC4 carries N-terminal sequences that are conserved across distantly related CC-NLRs. (**A**) Schematic representation of the different NLR domains used in TRIBE-MCL and phylogenetic analyses. (**B**) Distribution of plant NLRs across N-terminal domain tribes. The colour codes are: orange for dicot NLRs and blue for monocot NLRs. (**C**) NLRs from the same N-terminal tribe are dispersed across NLR phylogeny. The phylogenetic tree was generated in MEGA7 by the neighbour-joining method using the NB-ARC domain sequences of 988 CC-NLRs identified from *N. benthamiana*, tomato, sugar beet, Arabidopsis, rice and barley. Tribe 1 to Tribe 4 members are marked with different colours as indicated in each panel. Red arrow heads indicate bootstrap support > 0.7 and is shown for the relevant nodes. The scale bars indicate the evolutionary distance in amino acid substitution per site. The full phylogenetic tree can be found in Figure supplement 8.

We performed phylogenetic analyses of NLR proteins using the NB-ARC domain because it is the only conserved domain that produces reasonably good global alignments and can inform evolutionary relationships between all members of this family. We mapped individual NLR proteins grouped in Tribe-MCL N-terminal tribes onto a phylogenetic tree based on the NB-ARC domain (Figure 3C). These analyses revealed that the clustering of NLRs into the N-terminal tribes does not always match the NB-ARC phylogenetic clades (Figure 3C). In particular, NLRs in Tribe 1 and Tribe 2 often mapped to distinct well-supported clades scattered throughout the NB-ARC phylogenetic tree. We conclude that there are N-terminal domain sequences that have remained conserved over evolutionary time across distantly related CC-NLRs.

### NRC4 and ZAR1 share the N-terminal MADA motif

Next, we investigated whether N-terminal domains of CC-NLRs carry specific sequence motifs. We used MEME (Multiple EM for Motif Elicitation) (Bailey and Elkan, 1994) to identify conserved patterns in each of the N-terminal domain tribes. MEME revealed several conserved sequence patterns in each of the four largest tribes (Figure supplement 2). In particular, a motif that is conserved at the N terminus of 87 of 102 proteins from Tribe 2 overlapped with the N-terminal 29 amino acids of NRC4 we identified as sufficient to cause cell death (Figure supplement 2). Remarkably, the conserved sequence pattern of this very N-terminal motif matched the ZAR1 α1 helix that undergoes a conformational switch during activation of the ZAR1 resistosome (Wang et al., 2019b) (Figure 4A and 4B). In fact, 8 of the first 17 amino acids of ZAR1 are invariant in NRC4, and the majority of the amino acid polymorphisms between ZAR1 and NRC4 in the α1 helix region are conservative (Figure 4A). We conclude that NRC4, ZAR1 and numerous other CC-NLRs share a conserved N-terminal motif. We coined this sequence “MADA motif” based on the deduced 21 amino acid consensus sequence MADAxVSFxVxKLxxLLxxEx (Figure 4C).

**Figure 4.**
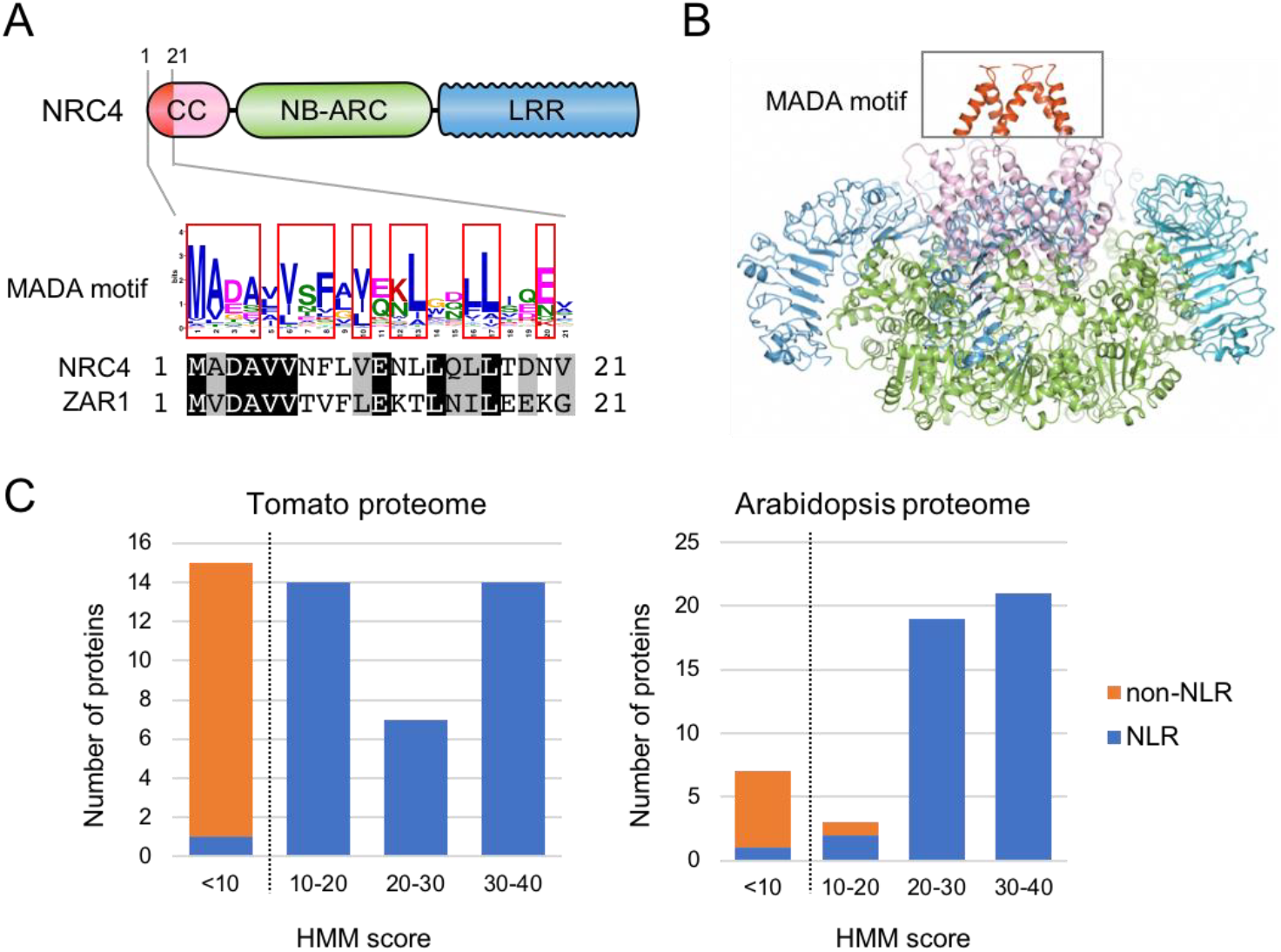
The MADA motif is a conserved unit at the very N-terminus of NRC4 and ZAR1. (**A**) Schematic representation of a classical CC-NLR protein highlighting the position of the MADA motif. Consensus sequence pattern of the MADA motif identified by MEME along with an alignment of NRC4 and ZAR1. Red boxes refer to residues conserved over 45% in Tribe 2 NLRs. (**B**) A structure homology model of NRC4 based on ZAR1 resistosome illustrating the position of the MADA motif. Each of the modelled five monomers is illustrated in cartoon representation. The colour code is: red for the MADA motif. The grey box highlights the N-terminal α helices, which contain the MADA motif. (**C**) Distribution of the MADA motif in tomato (left) and Arabidopsis (right) proteomes following HMMER searches with the MADA motif HMM. The number of proteins in each HMM score bin is shown. NLR and non-NLR proteins are shown in orange and blue, respectively. The dashed line indicates the cut-off used to define the most robust MADA-CC-NLR. NLRs with scores <10.0 were classified as MADA-like NLRs (MADAL-NLRs).

### The MADA motif is primarily found in NLR proteins

We built a Hidden Markov Model (HMM) from a sequence alignment of the MADA motif of 87 NLR proteins from Tribe 2 (Supplementary Table 2). To determine whether the MADA motif is primarily found among proteins annotated as NLRs, we used the HMMER software (Eddy, 1998) to query the Arabidopsis and tomato proteomes using the MADA motif HMM. HMMER searches revealed that the MADA motif is mainly found in NLR proteins compared with non-NLR proteins (Figure 4D; Supplementary Table 3). An HMM score cut-off of 10.0 clearly distinguishes NLR proteins from others with 100% (35 out of 35) tomato proteins and 97.7% (42 out of 43) Arabidopsis proteins scoring over 10.0 being annotated as NLRs (Figure 4D; Figure supplement 3). We conclude that the MADA motif is a sequence signature of a subset of NLR proteins and that a HMMER cut-off score of 10.0 is most optimal for high confidence searches of MADA containing CC-NLR proteins (MADA-CC-NLRs).

### MADA-like sequences occur in the N-termini of about 20% of dicot and monocot CC-NLRs

To what extent does the MADA motif occur in plant NLRomes? We re-screened the CC-NLR database using HMMER and identified 106 hits (10.7%) over the cut-off score of 10.0 (Figure 5A and 5B; Figure supplement 4A; Supplementary Table 4). We also noted that another 132 NLRs were positive but with a score lower than 10.0, and we tentatively termed these hits as MADA-like CC-NLRs (MADAL-CC-NLRs) (Figure 5B; Figure supplement 4A; Supplementary Table 4). Most of the MADA hits are from dicot plant species whereas MADA-like NLRs are primarily from monocots possibly reflecting a bias in our HMM profile which was built from the dicot enriched Tribe 2 (Figure 5B; Figure supplement 4B). Indeed, the majority of MADA hits (85 out of 106) were from Tribe 2, which includes NRC4 and ZAR1, but some MADA hits were also from other Tribes, notably the rice helper NLR Pik-2 from Tribe 1 (HMM score = 10.5) (Figure 5B; Figure supplement 4C). MADAL-CC-NLRs are mainly from Tribe 1 and Tribe 4 and include the monocot proteins MLA10 and Sr33, as well as Arabidopsis RPM1 (Figure 5B; Figure supplement 4C).

**Figure 5.**
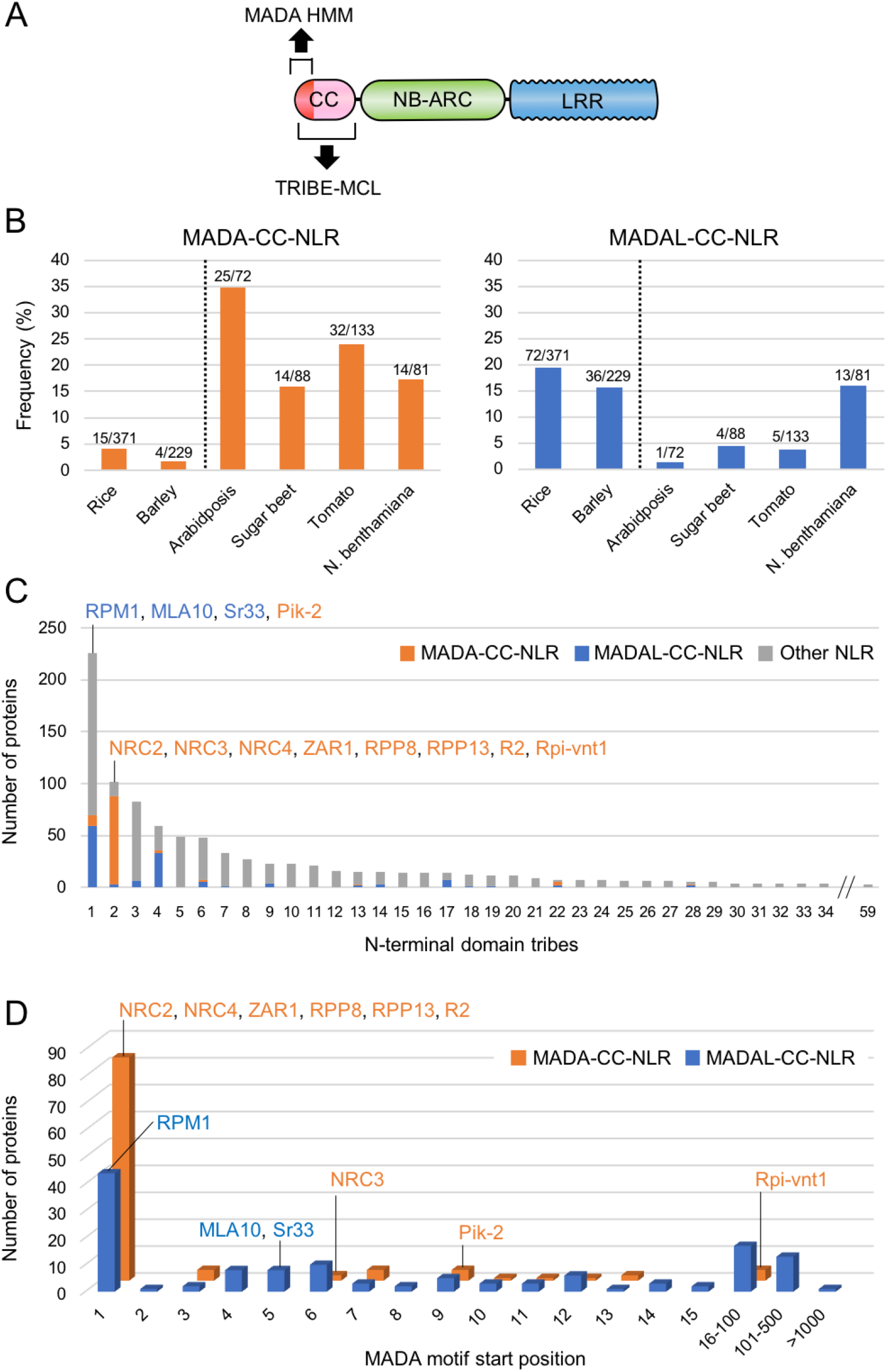
The MADA motif is conserved in ~20% of CC-NLRs. (**A**) Schematic representation of a classical CC-NLR protein highlighting the regions used for HMMER searches (MADA-HMM) and for TRIBE-MCL. (**B**) Occurrence of MADA/MADAL-CC-NLRs in representative species of monocots and dicots. The frequency of MADA/MADAL-CC-NLRs for each plant species was calculated as a percentage of all predicted CC-NLR proteins. (**C**) Occurrence of MADA/MADAL-CC-NLRs in N-terminal domain tribes of CC-NLRs. (**D**) Position distribution of MADA/MADA-like motif relative to the start codon position among the identified 106 MADA-CC-NLRs and 132 MADAL-CC-NLRs. The colour codes are: orange for MADA-CC-NLRs, blue for MADAL-CC-NLRs and grey for other NLRs.

Given that the MADA sequence is at the very N-terminus of ZAR1 and NRC4, and that the N-terminal position of the ZAR1 α1 helix is critical for its function based on the model of Wang et al. (2019b), we checked the positional distribution of predicted MADA and MADA-like motifs (Figure 5C). The majority of the predicted MADA and MADA-like motifs (203 out of 238, 85%) occurred at the very beginning of the NLR protein. However, 4 of 106 of the predicted MADA and 31 of 132 MADA-like NLRs have N-terminal extensions over 15 amino acids prior to the motifs (Figure 5C). For example, the MADA motif is located at position 52 to 72 amino acids in the potato NLR Rpi-vnt1. Whether these exceptions reflect misannotated gene models or genuinely distinct motif sequences remains to be determined.

In summary, our bioinformatic analyses revealed that 203 out 988 (20.5%) of the CC-NLRs of six representative dicot and monocot species contain a MADA or MADA-like motif at their very N-termini. These MADA sequences have noticeable similarity to NRC4 and ZAR1.

### All NRC-dependent sensor NLRs (NRC-S) lack the MADA motif

NB-ARC domain phylogenetic trees revealed that the NRC superclade is divided into the NRC clade (NRC-H) and a larger clade that includes all known NRC-dependent sensor NLRs (NRC-S) (Wu et al., 2017). We noted that even though the NRC-H and NRC-S are sister clades based on NB-ARC phylogenetic analyses, they grouped into distinct N-terminal domain tribes in the Tribe-MCL analyses (Figure 6A). Whereas all NRC-H mapped to Tribe 2, NRC-S clustered into 8 different tribes (Figure 6A). This pattern indicates that unlike the NRCs, the N-terminal sequences of their NRC-S mates have diversified throughout evolutionary time.

**Figure 6.**
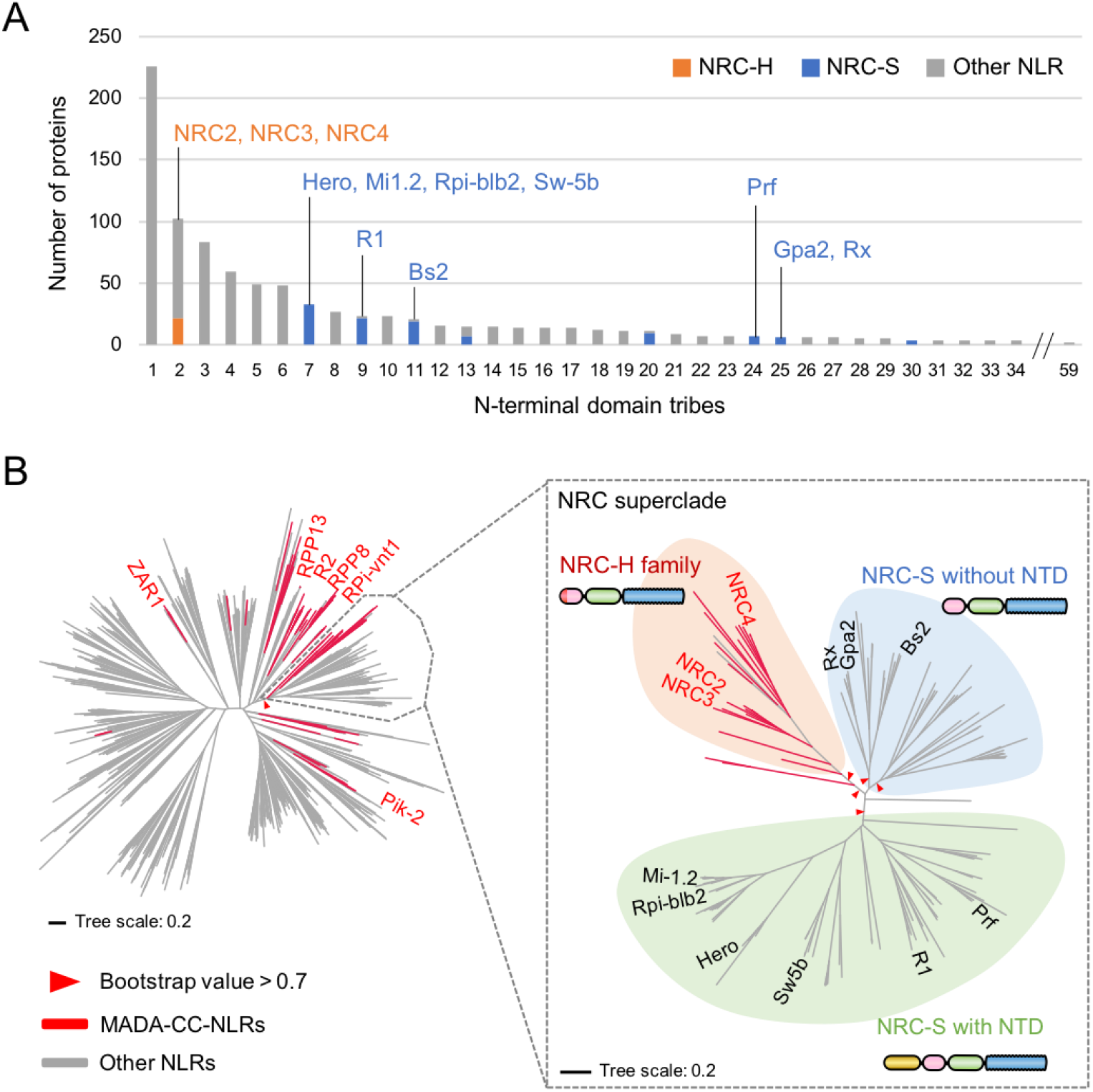
NRC-dependent sensors (NRC-S) do not have the MADA motif. (**A**) Distribution of NRCs (NRC-H) and NRC-dependent sensors (NRC-S) across N-terminal domain tribes of CC-NLRs. Individual NLR members of the NRC superclade were classified based on phylogenetic analysis as described in Figure supplement 8. The colour codes are: orange for the NRCs (NRC-H), blue for the NRC-sensors (NRC-S) and grey for other NLRs. (**B**) NRC-dependent sensors (NRC-S) do not contain the MADA motif. The phylogenetic tree of the 988 CC-NLRs described in Figure 3C is shown in the left panel with the NRC superclade marked by the grey lines. The NRC superclade phylogenetic tree is shown on the right panel and highlights the well-supported subclades NRC-H and the expanded NRC-S. The NRC-S clade is divided into NLRs that lack an N-terminal extension domain (NTD) prior to their CC domain and those that carry an NTD. MADA-CC-NLRs are highlighted in red in both trees. Red arrowheads mark bootstrap supports > 0.7 in relevant nodes. The scale bars indicate the evolutionary distance in amino acid substitution per site. The full phylogenetic tree can be found in Figure supplement 8. Schematic representation of domain architecture of the depicted classes of NLR protein is also shown similar to the other figures but with the ~600 amino acid NTD shown in yellow.

Next, we mapped the occurrence of the MADA motif onto the NB-ARC phylogenetic tree and noted that the distribution of the MADA motif was uneven across the NRC superclade despite their phylogenetic relationship (Figure 6B). Whereas 20 out of 22 NRC-H have a predicted MADA motif at their N-termini, none of the 117 examined NRC-S were predicted as MADA-CC-NLR in the HMMER search (Figure 6B; Supplementary Table 5). In fact, 65 of 117 NRC-S, including the well know disease resistance proteins R1, Prf, Sw5b, Hero, Rpi-blb2 and Mi-1.2, have N-terminal extensions of ~600 amino acids, or more in the case of Prf, prior to their predicted CC domains (Figure 6B). These findings indicate the CC domains of NRCs and their NRC-S mates have experienced distinct evolutionary trajectories even though these NLR proteins share a common evolutionary origin.

### MADA motif residues are required for NRC4 to trigger cell death

To experimentally validate our bioinformatic analyses, we performed site directed mutagenesis to determine the degree to which the MADA motif is required for the activity of NRC4. First, we followed up on the ZAR1 structure-function analyses of Wang et al. (2019b) who showed that three amino acids (phenylalanine 9 [F9], leucine 10 [L10] and leucine 14 [L14]) within the α1 helix/MADA motif are required for ZAR1-mediated cell death and bacterial resistance. We introduced a triple alanine substitution similar to the mutant of Wang et al. (2019b) into the autoactive NRC4^DV^ and found that this L9A/V10A/L14A mutation significantly reduced, but did not abolish, NRC4^DV^ cell death inducing activity (Figure 7A to 7C). Given that the MADA motif, particularly the mutated L9, V10 and L14 sites, is primarily composed of hydrophobic residues, we reasoned that substitutions with the negatively charged glutamic acid (E) would be more disruptive than hydrophobic alanine. Therefore, we substituted L9, V10 and L14 with glutamic acid, and observed that the L9E/L10E/L14E mutation resulted in a more severe disruption of the cell death activity of NRC4^DV^ compared to the triple alanine mutant (Figure 7A to 7C). Both of the NRC4^DV^ triple alanine and glutamic acid mutant proteins accumulated to similar levels as NRC4^DV^ when expressed in *N. benthamiana* leaves indicating that the observed loss-of-function phenotypes were not due to protein destabilization (Figure 7D).

**Figure 7.**
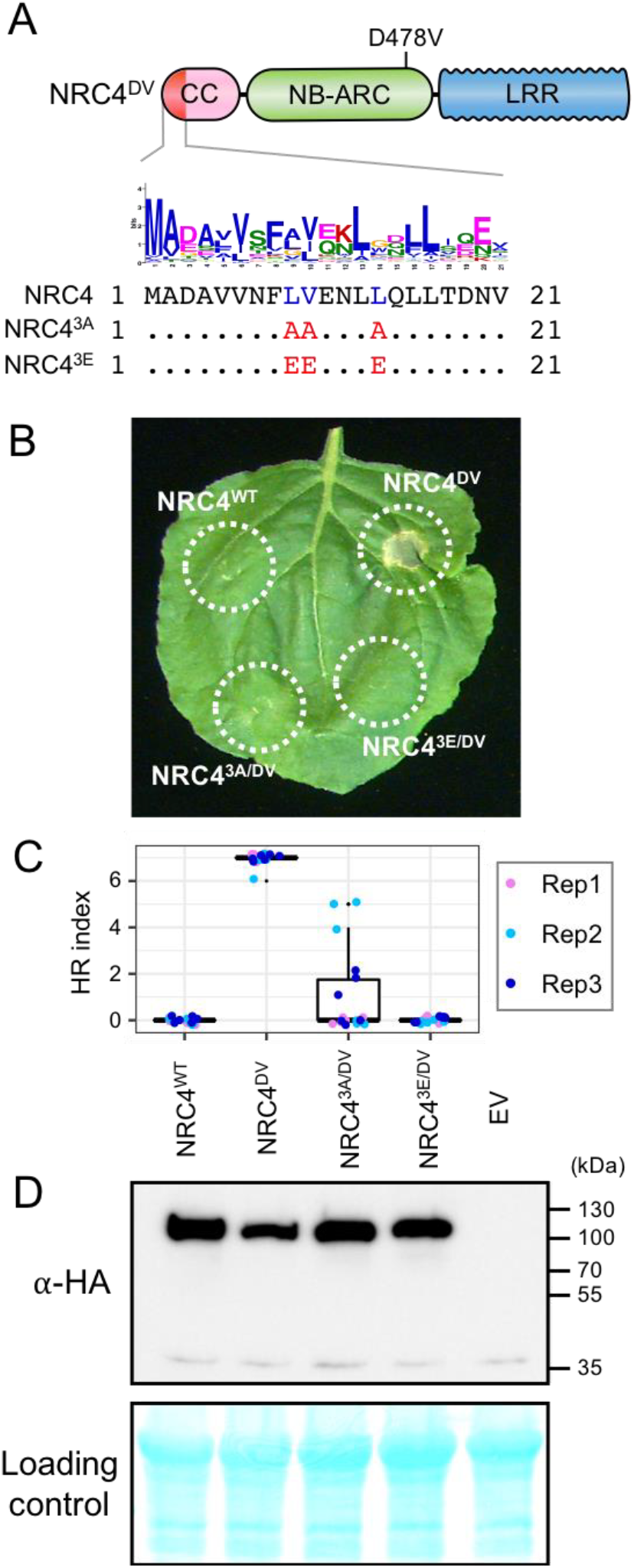
L9, V10 and L14 triple mutation impairs cell death activity of autoimmune NRC4^DV^. (**A**) Schematic representation of NRC4 and the mutated sites in the MADA motif. Mutated sites and substituted residues are shown as red characters in the NRC4 sequence alignment. (**B**) Cell death observed in *N. benthamiana* after expression of NRC4 mutants. *N. benthamiana* leaf panels expressing NRC4^WT^-6xHA, NRC4^DV^-6xHA, NRC4^3A/DV^-6xHA and NRC4^3E/DV^-6xHA were photographed at 5 days after agroinfiltration. (**C**) Box plots showing cell death intensity scored as an HR index based on three independent experiments. (**D**) *In planta* accumulation of the NRC4 variants. For anti-HA immunoblots of NRC4 and the mutant proteins, total proteins were prepared from *N. benthamiana* leaves at 1 day after agroinfiltration. Empty vector control is described as EV. Equal loading was checked with Reversible Protein Stain Kit (Thermo Fisher).

Next, we performed single alanine and glutamic acid mutant scans to reveal which other residues in the MADA motif are required for NRC4-mediated cell death. None of the tested single alanine-substituted mutants affected the cell death response of NRC4^DV^ (Figure supplement 5). In contrast, single glutamic acid mutations L9E, L13E and L17E essentially abolished the cell death activity of NRC4^DV^ without affecting the stability of the mutant proteins (Figure 8). Therefore, we determined that the L9, L13, and L17 residues in the MADA motif are critical for cell death induction by NRC4.

**Figure 8.**
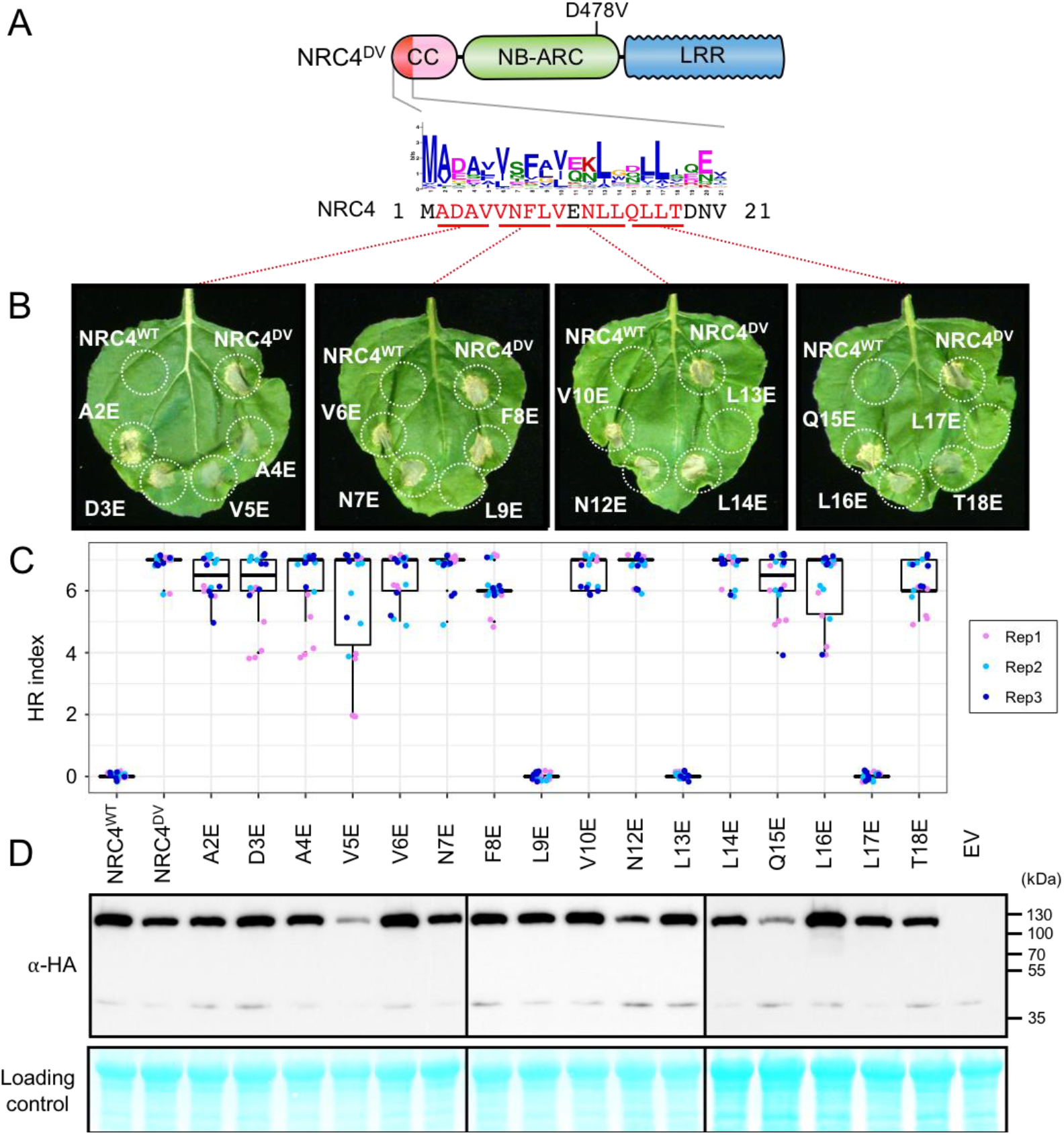
L9E, L13E and L17E single mutations impair cell death activity of autoimmune NRC4^DV^. (**A**) Schematic representation of NRC4 and the glutamic acid (E) mutant scan of the MADA motif. Mutated sites are shown as red characters in the NRC4 sequence. (**B**) Cell death observed in *N. benthamiana* after expression of NRC4 mutants. *N. benthamiana* leaf panels expressing NRC4^WT^-6xHA, NRC4^DV^-6xHA and the corresponding E mutants were photographed at 5 days after agroinfiltration. (**C**) Box plots showing cell death intensity scored as an HR index based on three independent experiments. (**D**) *In planta* accumulation of the NRC4 variants. Immunoblot analysis was done as described in Figure 7D.

Finally, we mapped L9, L13 and L17 onto a homology model of the CC domain of NRC4 produced based on the ZAR1 resistosome structure of Wang et al. (2019b) (Figure supplement 6). All three residues mapped to the outer surface of the funnel-shaped structure formed by the α1 helices similar to the previously identified residues in positions 9, 10 and 14. These results indicate that the outer surface of the funnel-shaped structure formed by N-terminal helices is critical not only for the function of ZAR1 but also for the activity of another MADA-CC-NLR.

### The α1 helix of Arabidopsis ZAR1 and the N-termini of other MADA-CC-NLRs can functionally replace the N-terminus of NRC4

Our observation that the ZAR1 α1 helix has sequence similarity to the N-terminus of NRC4 prompted us to determine whether this sequence is functionally conserved between these two proteins. To test this hypothesis, we swapped the first 17 amino acids of NRC4^DV^ with the equivalent region of ZAR1 (Figure 9A and 9B). The resulting ZAR1_1-17_-NRC4 chimeric protein can still trigger cell death in *N. benthamiana* leaves indicating that the MADA/α1 helix sequence is functionally equivalent between these two NLR proteins (Figure 9C).

**Figure 9.**
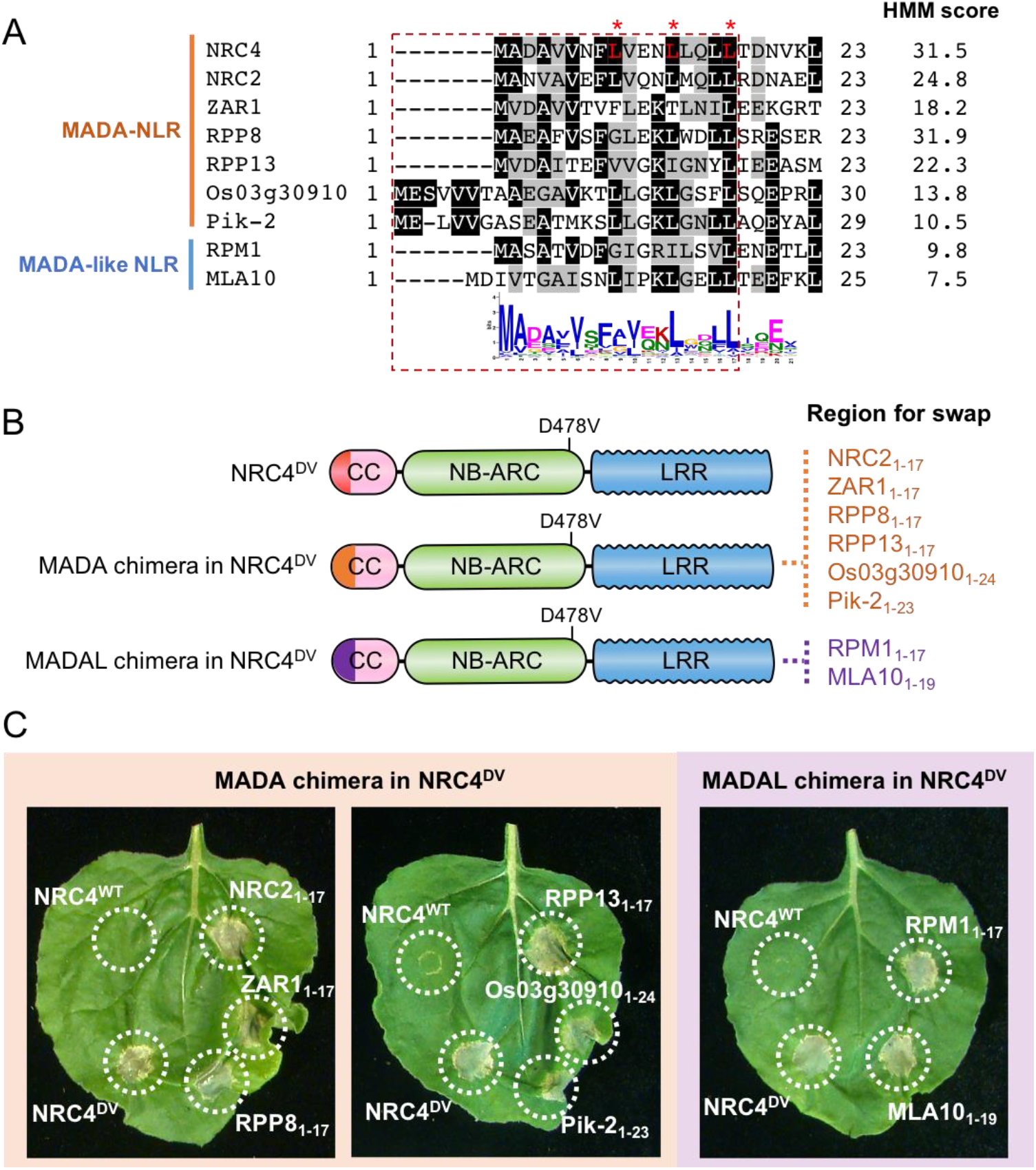
First 17 amino acids of NRC4 can be functionally replaced by the N-terminus of other MADA/MADALCC-NLRs. **(A)** Alignment of the N-terminal region of the MADA/MADAL-CC-NLRs. Key residues for cell death activity identified in Figure 8 are marked as red characters with asterisks in the sequence alignment. Each HMM score is indicated. **(B)** Schematic representation of NRC4 MADA motif chimeras with MADA/MADAL motifs from other CC-NLRs. The first 17 amino acid region of other MADA-CC-NLR (orange) or MADAL-CC-NLR (purple) was swapped into NRC4^DV^, resulting in the NRC4 chimeras with MADA/MADAL motifs originated from other NLRs. **(C)** Cell death phenotypes induced by the NRC4 chimeras. NRC4^WT^-6xHA, NRC4^DV^-6xHA and the chimeras were expressed in *N. benthamiana* leaves. Photographs were taken at 5 days after agroinfiltration.

Next, we swapped the same 17 amino acids of NRC4 with the matching sequences of the MADA-CC-NLRs NRC2 from *N. benthamiana*, RPP8 and RPP13 from Arabidopsis, and Pik-2 and Os03g30910.1 from rice, all of which gave HMMER scores >10.0 and ranging from 31.9 to 10.5 (Figure 9A and 9B). All of the assayed chimeric NRC4^DV^ proteins retained the capacity to trigger cell death in *N. benthamiana* leaves (Figure 9C). We further determined whether the N-termini of MADAL-CC-NLRs Arabidopsis RPM1 and barley MLA10, which yielded respective HMMER scores of 9.8 and 7.5, could also replace the first 17 amino acids of NRC4^DV^ (Figure 9A and 9B). Both NRC4^DV^ chimeras retained the capacity to trigger cell death indicating that these MADA-like sequences are functionally analogous to the NRC4 N-terminus (Figure 9C). Taken together, these results indicate that the MADA motif is functionally conserved even between distantly related NLRs from dicots and monocots.

### ZAR1-NRC4 chimeric protein retains the capacity to confer Rpi-blb2-mediated resistance against the late blight pathogen *Phytophthora infestans*

We investigated whether the MADA motif of NRC4 is required for disease resistance against the oomycete pathogen *Phytophthora infestans*. One of the NRC4-dependent sensor NLRs is Rpi-blb2, an NRC-S protein from *Solanum bulbocastanum* that confers resistance to *P. infestans* carrying the matching effector AVRblb2 (van der Vossen et al., 2003; Oh et al., 2009). For this purpose, we set up a genetic complementation assay in which NRC4 is co-expressed with Rpi-blb2 in leaves of the *N. benthamiana nrc4a/b* mutant prior to inoculation with the *P. infestans* strain 88069 (Wu et al., 2017), that carries AVRblb2 (Figure 10A). Unlike wild-type NRC4, the NRC4 L9A/V10A/L14A and L9E mutants failed to rescue the resistance to *P. infestans* in the *N. benthamiana nrc4a/b* mutant, indicating that MADA motif mutations not only impair HR cell death as shown above but also affect disease resistance against an oomycete pathogen (Figure 10B). We conducted similar complementation assays with the ZAR1_1-17_-NRC4 chimera in which the first 17 amino acids of NRC4 were swapped with the equivalent region of ZAR1, and found that ZAR1_1-17_-NRC4 complemented the *nrc4a/b N. benthamiana* mutant to a similar degree as wild-type NRC4 (Figure 10B). These experiments further confirm that the α1 helix/MADA motif of Arabidopsis ZAR1 is functionally equivalent to the N-terminus of NRC4, and that the chimeric ZAR1_1-17_-NRC4 is not only able to trigger HR cell death but also retains its capacity to function with its NRC-S mate Rpi-blb2 and confer resistance to *P. infestans*.

**Figure 10.**
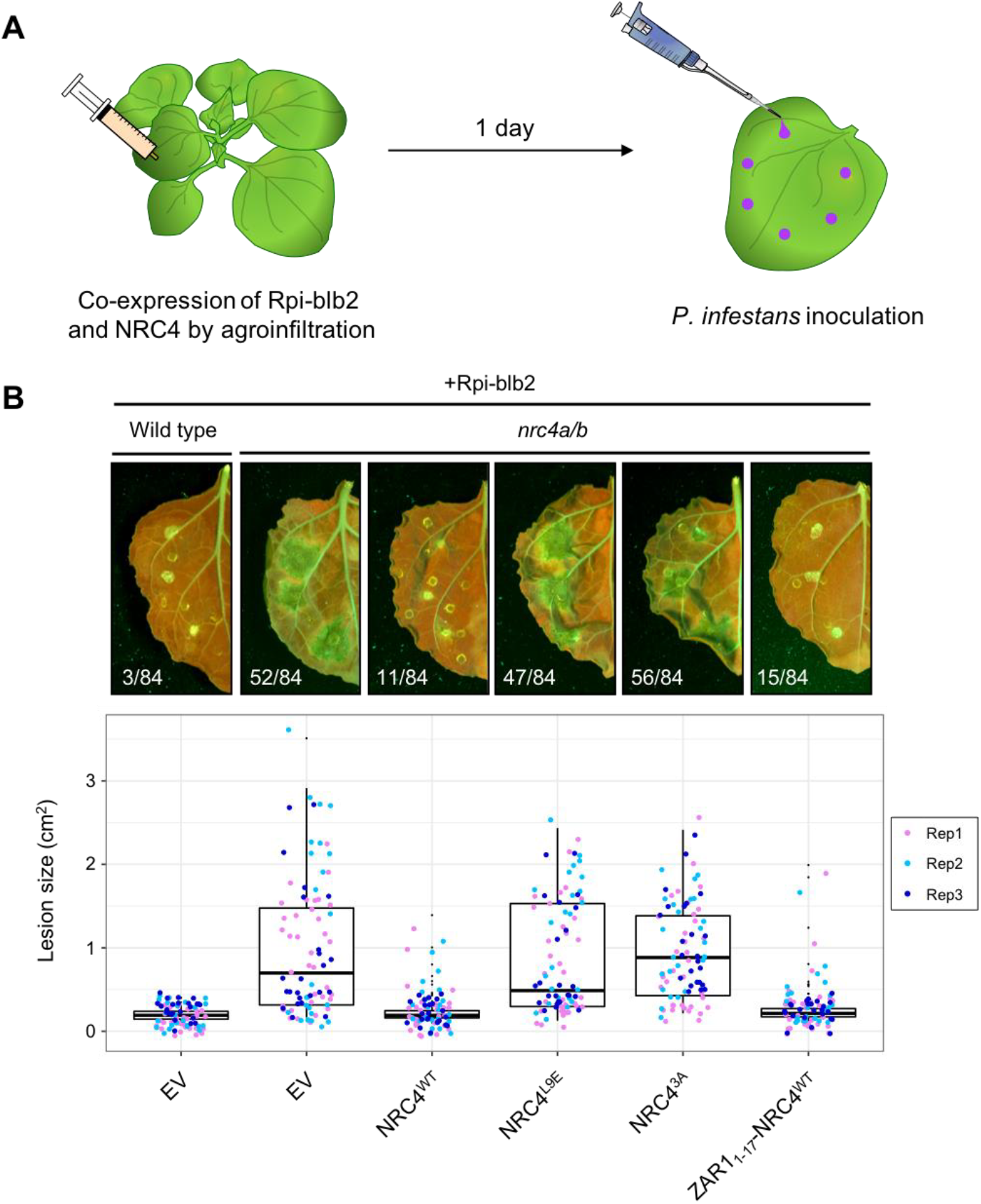
The chimeric protein ZAR1_1-17_-NRC4 complements NRC4 function in Rpi-blb2-mediated resistance. **(A)**Schematic representation of NRC4 complementation assay for Rpi-blb2-mediated resistance. Wild-type and the variants of NRC4 were co-expressed with RFP-Rpiblb2 in wild-type or *nrc4a/b N. benthamiana* leaves by agroinfiltration. The leaves were inoculated with droplets of zoospore suspension from *P. infestans* strain 88069 at 1 day after the agroinfiltration. The syringe and pipet are not drawn to scale. (**B**) Disease and resistance phenotypes on NRC4/Rpi-blb2-expressed leaves. Images were taken under UV light at 7 days post inoculation. The lesion size (bottom panel) was measured in Fiji. Experiments were repeated three times with totally 84 inoculation site each. The numbers on the photographs indicate the sum of spreading lesions/total inoculation sites from the three replicates.

## Discussion

This study stems from a random truncation screen of the CC-NLR NRC4, which revealed that the very N-terminus of this protein is sufficient to carry out the HR cell death activity of the full-length protein. It turned out that this region is defined by a consensus sequence—the MADA motif—that occurs in about one fifth of plant CC-NLRs including Arabidopsis ZAR1. The MADA motif covers most of the functionally essential α1 helix of ZAR1 that undergoes a conformational switch during activation of the ZAR1 resistosome (Wang et al., 2019b). Our finding that the ZAR1 α1 helix/MADA motif can functionally replace its matching region in NRC4 indicates that the ZAR1 “death switch” mechanism may apply to NRCs and other MADA-CC-NLRs from dicot and monocot plant species.

We recently proposed that NLRs may have evolved from multifunctional singleton receptors to functionally specialized and diversified receptor pairs and networks (Adachi et al., 2019a). In this study, a striking finding from the computational analyses is that all NRC-S lack the MADA motif even though they are more closely related to NRC-H than to ZAR1 and other MADA-CC-NLRs in the NB-ARC phylogenetic tree (Figure 6). These observations led us to draw the evolutionary model of Figure 11. In this model, we propose that MADA-type sequences have emerged early in the evolution of CC-NLRs and have remained conserved from singletons to helpers in NLR pair and network throughout evolution. In sharp contrast, MADA sequences appear to have degenerated over time in sensor CC-NLRs as these proteins specialized in pathogen detection and lost the capacity to execute the immune response without their helper mates. Consistent with this view, NRCs are known to be more highly conserved than their NRC-S partners within the Solanaceae (Wu et al. 2017; Stam et al., 2019). Future analyses will determine whether MADA-CC-NLRs are generally more evolutionarily constrained than non-MADA containing NLRs.

**Figure 11.**
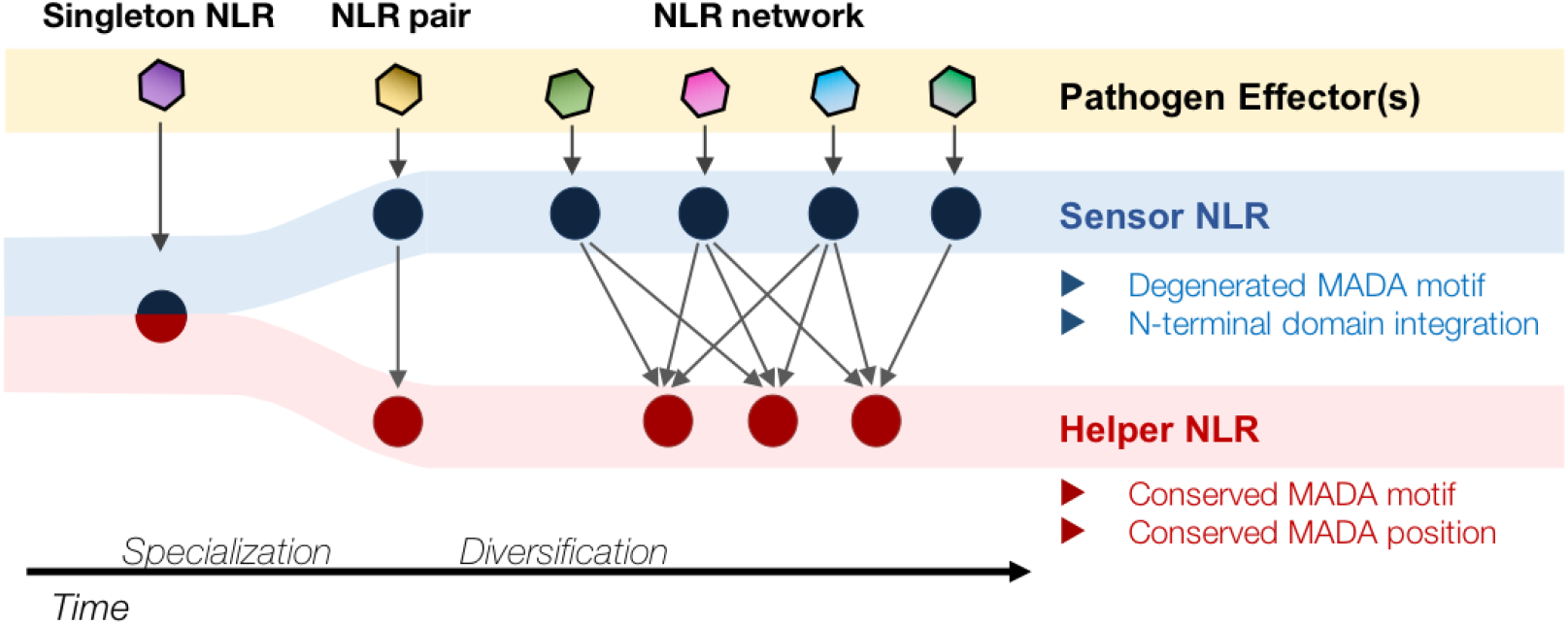
Evolution of NLRs from singletons to networks. We propose that the N-terminal MADA motif/α1 helix has emerged early in the evolution of CC-NLRs and has remained constrained throughout time as singletons evolved from multifunctional proteins into specialized paired and networked NLR helpers. In contrast, the MADA motif/α1 helix has degenerated in sensor CC-NLRs as they rely on their NLR helper mates for executing the immune response (“use-it-or-lose-it” model of evolution). In addition, some sensor NLRs, such as a large subset of NRC-S proteins, have acquired N-terminal domains (NTDs)—prior to their CC domains—that function in pathogen detection. Such NTDs would preclude a free N-terminal α1 helix, which would be incompatible with the current model of ZAR1 resistosome activation.

Consistent with our model, the CC domain of some NRC-S, such as Sw5b and Rx, is dispensable for activation of HR cell death (De Oliveira et al., 2016; Zhu et al., 2017; Rairdan et al., 2008). In addition, about half of the NRC-S proteins have acquired N-terminal extensions (N-terminal domains) before their CC domain, which would preclude a free N-terminal α1 helix essential for a ZAR1 type “death switch” mechanism (Figure 6). In fact, the N-terminal domains of Prf and Sw5b function as baits that sense pathogen effectors, suggesting functional analogy to integrated effector detection motifs found in some NLRs, and are not known to be involved in executing the immune response (Saur et al., 2015; Li et al., 2019). Here, we hypothesize that the CC domains of these and other sensor NLRs have extensively diversified over evolutionary time and lost the capacity to function as HR cell death executors. This could be a consequence of relaxed selection given that these proteins rely on their MADA-CC-NLR partners to execute the immune response as discussed above. Additional structure-function experiments will be needed to determine the extent to which this “use-it-or-lose-it” evolutionary model applies to the sensor sub-class of NLR immune receptors.

Already, our evolutionary model appears to be consistent with some paired NLR configurations in addition to the NRC-H/NRC-S network. One example is rice Pik-1 and Pik-2, which are a well-established NLR pair that detects the AVRPik effector of the rice blast fungus *M. oryzae* (Maqbool et al., 2015; Białas et al., 2018). AVRPik binding to the integrated heavy metal associated (HMA) domain of Pik-1 results in HR cell death and blast fungus resistance only in the presence of its helper Pik-2 protein (Maqbool et al., 2015). In our computational analyses only Pik-2 was detected to carry an N-terminal MADA motif (Figure 5, HMM score = 10.5) even though the CC domains of both proteins grouped into Tribe 1 (Figure 3). The Pik-2 MADA motif could substitute for the N-terminus of NRC4 in our cell death assays despite having 6 additional amino-acids at its N-terminus (Figure 9). These results are consistent with our Figure 11 model and imply that the helper NLR Pik-2 may execute HR cell death via its N-terminal MADA motif whereas its paired sensor NLR Pik-1 does not have the capacity to carry this activity on its own.

In addition to ZAR1, RPP8 is another Arabidopsis MADA-CC-NLR with high similarity to the N-terminus of NRC4 with 9 invariant amino acids out of 17 (53%; HMMER score = 31.9). This RPP8 MADA motif could substitute for the N-terminus of NRC4 indicating that it is functional (Figure 9). In Arabidopsis, RPP8 (AT5G43470) and its paralogs occur at dynamic genetic loci that exhibit frequent sequence exchanges as deduced from comparative genomic analyses (Kuang et al., 2008). Four of the five RPP8 paralogs in the Arabidopsis ecotype Col-0 were deemed to have a MADA motif based on our HMMER searches, whereas a fifth paralog LOV1 (AT1G10920) was negative (Supplementary Table 6). LOV1 confers sensitivity to the victorin effector produced by the necrotrophic fungus *Cochliobolus victoriae* by binding the defense-associated thioredoxin TRX-h5 when it is complexed with victorin (Lorang et al., 2012). Interestingly, LOV1 binds TRX-h5 via its CC domain indicating that this region has evolved a pathogen sensor activity in this NLR protein (Lorang et al., 2012). How the sensor activity of the CC domain of LOV1 relates to the absence of a detectable MADA motif and whether this protein relies on other MADA-CC-NLRs to execute the cell death response are unanswered questions that are raised by these observations.

In activated ZAR1 resistosome, a funnel-shaped structure formed by five α1 helices is thought to directly execute HR cell death by forming a toxin-like pore in the plasma membrane (Wang et al., 2019b). To what extent do activated MADA-CC-NLRs function according to this ZAR1 model? Structure informed mutagenesis of ZAR1 revealed that F9, L10 and L14 on the outer surface of the funnel-shaped structure are required for immunity (Wang et al., 2019b). Here, our Ala and Glu scans of the MADA motif revealed that the NRC4 L9, L13 and L17 residues are essential for HR cell death activity. All three residues mapped to the outer surface of NRC4 α1 helices as predicted from a homology model based on the ZAR1 resistosome (Figure Supplement 6). These residues may be important for insertion of the funnel-shaped structure into the plasma membrane or interaction with downstream components that remain to be identified. As discussed by Wang et al. (2019b), the interior space of the funnel structure is also important because the ZAR1 double mutant E11A/E18A is impaired in cell death and disease resistance activities. However, in our Glu mutant scan, we failed to observe a reduction in HR cell death activities with single site mutants in these residues or other amino acids that are predicted to line up the interior space of the funnel-shaped structure. Whether or not this reflects genuine biological differences between ZAR1 and NRC4 remains to be studied.

A subset of CC-NLRs of the RPW8/HR family of atypical resistance proteins have a distinct type of coiled-coil domain known as CC_R_ (Barragan et al., 2019; Li et al., 2019). We failed to detect any MADA type sequences in these CC_R_-NLR proteins. Indeed, the CC_R_ domain has similarity to mixed lineage kinase domain-like (MLKL) proteins and fungal HeLo/HELL domains, which form multi-helix bundles and act as membrane pore forming toxins (Barragan et al., 2019; Li et al., 2019; Mahdi et al., 2019). Whether the CC_R_ domains function as a distinct cell death inducing system in plants compared to MADA-CC-NLRs remains to be determined. Interestingly, Arabidopsis HR4, a member of the CC_R_-NLR family, interacts in an allele-specific manner with the genetically unlinked CC-NLR RPP7b to trigger autoimmunity in the absence of pathogens (Barragan et al., 2019). Recently, Li et al. (2019) showed that RPP7b forms higher-order complexes of six to seven subunits only when activated by the matching autoimmune HR4^Fei-0^ protein in a biochemical process reminiscent of activated ZAR1 resistosome (Li et al., 2019). In our HMMER searches, RPP7b and its four Arabidopsis paralogs were all classed as carrying the MADA motif (Supplementary Table 6, HMMER score = 31.1). Thus, findings by Li et al. (2019) directly link a MADA-CC-NLR to the formation of resistosome type structures consistent with our view that the ZAR1 model widely applies to other NLRs with the MADA α1 helix. It will be fascinating to determine whether or not RPP7b and HR4 are both capable of executing cell death, especially as two-component systems of NLR and HeLo/HELL proteins are common in fungi and mammals (Barragan et al., 2019).

Plant NLRs can be functionally categorized into singleton, sensor or helper NLRs based on their biological activities (Adachi et al., 2019a). However, it remains challenging to predict NLR functions from the wealth of unclassified NLRomes that are emerging from plant genome sequences. It has not escaped our attention that the discovery of the MADA motif as a signature of NLR singletons and helpers—but missing in sensor NLRs—enables the development of computational pipelines for predicting NLR networks from naïve plant genomes. Such *in silico* predictions can be tested by co-expression of paired NLRs in *N. benthamiana*. In addition, MADA motif predictions can be validated using our straightforward functional assay of swapping the NRC4 N-terminus, with the readouts consisting of both HR cell death (Figure 9) and resistance to *P*. *infestans* (Figure 10). Dissecting the NLR network architecture of plant species is not only useful for basic mechanistic studies but has also direct implications for breeding disease resistance into crop plants and reducing the autoimmune load of NLRs (Chae et al., 2016; Wu et al., 2018; Adachi et al., 2019a).

## Materials and Methods

### Plant growth conditions

Wild type and mutant *N. benthamiana* were propagated in a glasshouse and, for most experiments, were grown in a controlled growth chamber with temperature 22-25°C, humidity 45-65% and 16/8-h light/dark cycle.

### Generation of *N. benthamiana nrc4a/b* CRISPR/Cas9 mutants

Constructs for generating *NRC4* knockout *N. benthamiana* were assembled using the Golden Gate cloning method (Weber et al., 2011; Nekrasov et al., 2013; Belhaj et al., 2013). sgRNA4.1 and sgRNA4.2 were cloned under the control of the Arabidopsis (*Arabidopsis thaliana*) U6 promoter (AtU6pro) [pICSL90002, The Sainsbury Laboratory (TSL) SynBio] and assembled in pICH47751 (addgene no. 48002) and pICH47761 (addgene no. 48003), respectively as previously described (Belhaj et al., 2013). Primers sgNbNRC4.1_F (tgtggtctcaATTGAAAAACGGTACATACCGCAGgttttagagctagaaatagcaag), sgNbNRC4.2_F (tgtggtctcaATTGAGTCAGGAATCTTGCAGCTGgttttagagctagaaatagcaag) and sgRNA_R (tgtggtctcaAGCGTAATGCCAACTTTGTAC) were used to clone sgRNA4.1 and sgRNA4.2. pICSL11017::NOSpro::BAR (TSL SynBio), pICSL11021::35Spro::Cas9 (addgene no. 49771), pICH47751::AtU6p::sgRNA4.1, pICH47761::AtU6pro::sgRNA4.2, and the linker pICH41780 (addgene no. 48019) were assembled into the vector pICSL4723 (addgene no. 48015) as described (Weber et al., 2011) resulting in construct pICSL4723::BAR::Cas9::sgRNA4.1::sgRNA4.2 that was used for plant transformation. Transgenic *N. benthamiana* were generated by TSL Plant Transformation team as described before (Nekrasov et al., 2013).

### *N. benthamiana nrc4a/b* genotyping

Genomic DNA of selected T2 *N. benthamiana* transgenic plants *nrc4a/b*_9.1.3 and *nrc4a/b*_1.2.1 was extracted using DNeasy Plant DNA Extraction Kit (Qiagen). Primers NRC4_1_F (GGAAGTGCAAAGGGAGAGTT), NRC4_1_R (TCGCCTGAACCACAAACTTA), NRC4_2_F (GGCAAGAATTTTGGATGTGG) and NRC4_2_R (CGAGGAACCCTTTTTAGGCAG) were used in multiplex polymerase chain reaction (PCR) assays to amplify the region targeted by the two sgRNAs. Multiplex amplicon sequencing was performed by the Hi-Plex technique (Lyon et al., 2016). Sequence reads were aligned to the reference *N. benthamiana* draft genome Niben.genome.v0.4.4 [Sol Genomics Network (SGN), https://solgenomics.net/], and *NRC4a* (on scaffold Niben044Scf00002971) and *NRC4b* (on scaffold Niben044Scf00016103) were further analysed. T3 lines from the selected T2 plants were used for the experiments.

### Plasmid constructions

To generate NRC4_1-29_-YFP expression construct, NRC4_1-29_ coding sequence was amplified by Phusion High-Fidelity DNA Polymerase (Thermo Fisher), and the purified amplicon was directly used in Golden Gate assembly with pICH85281 [mannopine synthase promoter+Ω (MasΩpro), addgene no. 50272], pICSL50005 (YFP, TSL SynBio), pICSL60008 [Arabidopsis heat shock protein terminator (HSPter), TSL SynBio] into binary vector pICH47742 (addgene no. 48001). Primers used for NRC4_1-29_ coding sequences are listed in Supplementary Table 7.

To generate an autoactive mutant of *N. benthamiana* NRC4, the aspartic acid (D) in the MHD motif was substituted to valine (V) by site-directed mutagenesis using Phusion High-Fidelity DNA Polymerase (Thermo Fisher). pCR8::NRC4^WT^ (Wu et al., 2017) was used as a template. Primers NRC4_D478V_F (5’-Phos/ATGTTGCATCAGTTCTGCAAAAAGGAGGCT) and NRC4_D478V_R (5’-Phos/GACGTGAAGACGACATGTTTTTATTTGACC) were used for introducing the mutation in the PCR. The mutated NRC4 was verified by DNA sequencing of the obtained plasmid.

pCR8::NRC4^WT^ (Wu et al., 2017) or pCR8::NRC4^DV^ without its stop codon were used as a level 0 modules for the following Golden Gate cloning. NRC4^DV^-3xFLAG was generated by Golden Gate assembly with pICH51266 [35S promoter+Ω promoter, addgene no. 50267], pICSL50007 (3xFLAG, addgene no. 50308) and pICH41432 (octopine synthase terminator, addgene no. 50343) into binary vector pICH47732 (addgene no. 48000). NRC4^WT^-6xHA and NRC4^DV^-6xHA were generated by Golden Gate assembly with pICH85281 (MasΩpro), pICSL50009 (6xHA, addgene no. 50309), pICSL60008 (HSPter) into the binary vector pICH47742. NRC4^WT^-YFP and NRC4^DV^-YFP were generated by Golden Gate assembly with pICH85281 (MasΩpro), pICSL50005 (YFP), pICSL60008 (HSPter) into binary vector pICH47742. For free YFP expression construct, pAGM3212 (YFP, TSL SynBio) was assembled with pICH85281 (MasΩpro) and pICSL60008 (HSPter) into the binary vector pICH47742 by Golden Gate reaction.

To generate MADA motif mutants and chimeras of NRC4, the full-length sequence of NRC4^WT^ or NRC4^DV^ was amplified by Phusion High-Fidelity DNA Polymerase (Thermo Fisher) with the forward primers listed in Supplementary Table 7. Purified amplicons were cloned into a level 0 vector pAGM1287 (addgene no. 47996) by Golden Gate cloning. The level 0 plasmids were then used for Golden Gate assembly with pICH85281 (MasΩpro), pICSL50009 (6xHA) and pICSL60008 (HSPter) into the binary vector pICH47742.

### Mu-STOP *in vitro* transposition

To generate the Mu-STOP transposon (Poussu et al., 2005), entranceposon M1-KanR (Mutation Generation System Kit, Thermo Fisher) was used as a PCR template, and three translational stop signals were added to each transposon end by Phusion High-Fidelity DNA Polymerase and Mu-STOP primer (GGAAGATCTGATTGATTGAACGAAAAACGCGAAAGCGTTTC). The 3’ A overhang was then introduced to the Mu-STOP amplicon by DreamTaq DNA polymerase (Thermo Fisher), and the resulting Mu-STOP amplicon was cloned into pGEM-T Easy (Promega). Mu-STOP transposon was then released from pGEM::Mu-STOP by *Bgl*II digestion and purified by GeneJET Gel Extraction Kit (Thermo Fisher). 100 ng of the purified Mu-STOP transposon was mixed with 500 ng of the target plasmid, pICH47732::35SΩpro::NRC4^DV^-3xFLAG, and MuA transposase from the Mutation Generation System Kit (Thermo Fisher). The *in vitro* transposition reaction was performed according to the manufacturer’s procedure and carried out at 30°C for 6 hours.

The NRC4^DV^::Mu-STOP library was transformed into *Agrobacterium tumefaciens* Gv3101 by electroporation. Mu-STOP insertion sites were determined by colony PCR using DreamTaq DNA polymerase (Thermo Fisher) and PCR amplicon sequencing. For the PCR, we used a forward primer (GAACCCTGTGGTTGGCATGCACATAC) matching pICH47732 and a reverse primer (CAACGTGGCTTACTAGGATC) matching Mu-STOP transposon.

### Transient gene-expression and cell death assays

Transient expression of NRC wild-type and mutants, as well as other genes, in *N. benthamiana* were performed by agroinfiltration according to methods described by Bos et al. (2006). Briefly, four-weeks old *N. benthamiana* plants were infiltrated with *A. tumefaciens* strains carrying the binary expression plasmids. *A. tumefaciens* suspensions were prepared in infiltration buffer (10 mM MES, 10 mM MgCl_2_, and 150 μM acetosyringone, pH5.6) and were adjusted to OD_600_ = 0.5. For transient expression of NRC4^WT^-YFP, NRC4^DV^-YFP, NRC4_1-29_-YFP or free YFP, the *A. tumefaciens* suspensions (OD_600_ = 0.25) were mixed in a 1:1 ratio with an *A. tumefaciens* expressing p19, the suppressor of posttranscriptional gene silencing of *Tomato bushy stunt virus* that is known to enhance *in planta* protein expression (Lindbo, 2007). HR cell death phenotypes were scored according to the scale of Segretin et al. (2014) modified to range from 0 (no visible necrosis) to 7 (fully confluent necrosis).

### Protein immunoblotting

Protein samples were prepared from six discs (8 mm diameter) cut out of *N. benthamiana* leaves at 1 day after agroinfiltration and were homogenised in extraction buffer [10% glycerol, 25 mM Tris-HCl, pH 7.5, 1 mM EDTA, 150 mM NaCl, 2% (w/v) PVPP, 10 mM DTT, 1x protease inhibitor cocktail (SIGMA), 0.2% IGEPAL (SIGMA)]. The supernatant obtained after centrifugation at 12,000 x*g* for 10 min was used for SDS-PAGE. Immunoblotting was performed with HA-probe (F-7) HRP (Santa Cruz Biotech) or anti-GFP antibody (ab290, abcam) in a 1:5,000 dilution. Equal loading was checked by taking images of the stained PVDF membranes with Pierce Reversible Protein Stain Kit (#24585, Thermo Fisher).

### Bioinformatic and phylogenetic analyses

We used NLR-parser (Steuernagel et al., 2015) to identify NLR sequences from the protein databases of tomato (SGN, Tomato ITAG release 2.40), *N. benthamiana* (SGN, *N. benthamiana* Genome v0.4.4), Arabidopsis (https://www.araport.org/, Araport11), sugar beet (http://bvseq.molgen.mpg.de/index.shtml, RefBeet-1.2), rice (http://rice.plantbiology.msu.edu/, Rice Gene Models in Release 7) and barley (https://www.barleygenome.org.uk/, IBSC_v2). The obtained NLR sequences, from NLR-parser, were aligned using MAFFT v. 7 (Katoh and Standley, 2013), and the protein sequences that lacked the p-loop motif were discarded from the NLR dataset. The gaps in the alignments were deleted manually in MEGA7 (Kumar et al., 2016) and the NB-ARC domains were used for generating phylogenetic trees (Supplementary file 4). The neighbour-joining tree was made using MEGA7 with JTT model and bootstrap values based on 100 iterations (Figure supplement 7). We removed TIR-NLR clade members from the final database, and retained all CC-NLR sequences, including the CC_R_-NLR (RPW8-NLR), that possess N-terminal domains longer than 30 amino acids (988 protein sequences, Supplementary file 2).

The NB-ARC domain sequences from 988 proteins (Supplementary file 5) were used to construct the CC-NLR phylogenetic tree in Figure supplement 8. The neighbour-joining tree was constructed as described above.

For the tribe analyses, we extracted the N-terminal domain sequences, prior to NB-ARC domain, from the CC-NLR database (Supplementary file 3), and used the Tribe-MCL feature from Markov Cluster Algorithm (Enright et al., 2002) to cluster the sequences into tribes with BLASTP E-value cutoff < 10^−8^. NLRs in each tribe were subjected to motif searches using the MEME (Multiple EM for Motif Elicitation) v. 5.0.5 (Bailey and Elkan, 1994) with parameters “zero or one occurrence per sequence, top 4 motifs”, to detect consensus motifs conserved in >= 70% of input sequences.

We used the most N-terminal motif detected in Tribe 2 from the MEME analysis to construct a hidden Markov model (HMM) for the MADA motif. Sequences aligned to the MADA motif were extracted in Stockholm format and used in hmmbuild program implemented in HMMER v2.3.2 (Eddy, 1998). The HMM was then calibrated with hmmcalibrate. This MADA-HMM (Supplementary file 6) was used to search the CC-NLR database (Supplementary file 2) with the hmmsearch program (hmmsearch --max -o <outputfile> <hmmfile> <seqdb>). To estimate the false positive rate, hmmsearch program was applied to full Arabidopsis and tomato proteomes (Araport11 and ITAG3.2) with the MADA-HMM and the output is displayed in Supplementary Table 3 and discussed in the results section.

### Pathogen infection assays

*P. infestans* infection assays were performed by applying droplets of zoospore suspension on detached leaves as described previously (Song et al., 2009). Briefly, leaves of five-weeks old wild-type and *nrc4a/b N. benthamiana* plants were infiltrated with *A. tumefaciens* solutions, in which each *Agrobacterium* containing a plasmid expressing RFP::Rpi-blb2 (Wu et al., 2017) was mixed in a 1:1 ratio (OD_600_ = 0.5 for each strain) with *Agrobacterium* containing either the empty vector, wild type NRC4, or NRC4 variant. At 24 hours after agroinfiltration, the abaxial side of the leaves were inoculated with 10 µL zoospore suspension (100 zoospores/μL) of *P. infestans* strain 88069 prepared according to the methods reported by Song et al. (2009). The inoculated leaves were kept in a moist chamber at room temperature (21-24°C) for 7 days, and imaged under UV light for visualization of the lesions.

### Structure homology modelling

We used the cryo-EM structure of activated ZAR1 (Wang et al. 2019b) as template to generate a homology model of NRC4. The amino acid sequence of NRC4 was submitted to Protein Homology Recognition Engine V2.0 (Phyre2) for modelling (Kelley et a., 2015). The coordinates of ZAR1 structure (6J5T) were retrieved from the Protein Data Bank and assigned as modelling template by using Phyre2 Expert Mode. The resulting model of NRC4 comprised amino acids Val-5 to Glu-843 and was illustrated in CCP4MG software (McNicholas et al., 2011).

### Accession Numbers

The NRC4 sequences used in this study can be found in the Solanaceae Genomics Network (SGN) or GenBank/EMBL databases with the following accession numbers: NbNRC4 (NbNRC4, MK692737; NbNRC4a, NbS00002971; NbNRC4b, NbS00016103).

## Acknowledgements

We are thankful to several colleagues for discussions and ideas. We thank Matthew Smoker and other members of the TSL Plant Transformation facility as well as Mark Youles of TSL SynBio for invaluable technical support. We thank Kurt Lamour for advice on genotyping the CRISPR/Cas9 mutants. H.A. is funded by the Japan Society for the Promotion of Science (JSPS) and L.D. by a Marie Sklodowska-Curie Actions (MSCA) Fellowship. The Kamoun Lab is funded primarily from the Gatsby Charitable Foundation, Biotechnology and Biological Sciences Research Council (BBSRC, UK), and European Research Council (ERC; NGRB and BLASTOFF projects).

## Author contributions

H.A. and S.K. designed the research and wrote the paper; H.A., M.C., A.H., C.-H.W, L.D., T.S., C.D. and E.M. performed research; and T.O.B., A.M., J.W. and S.K. supervised research.

---

**Figure supplement 1.**
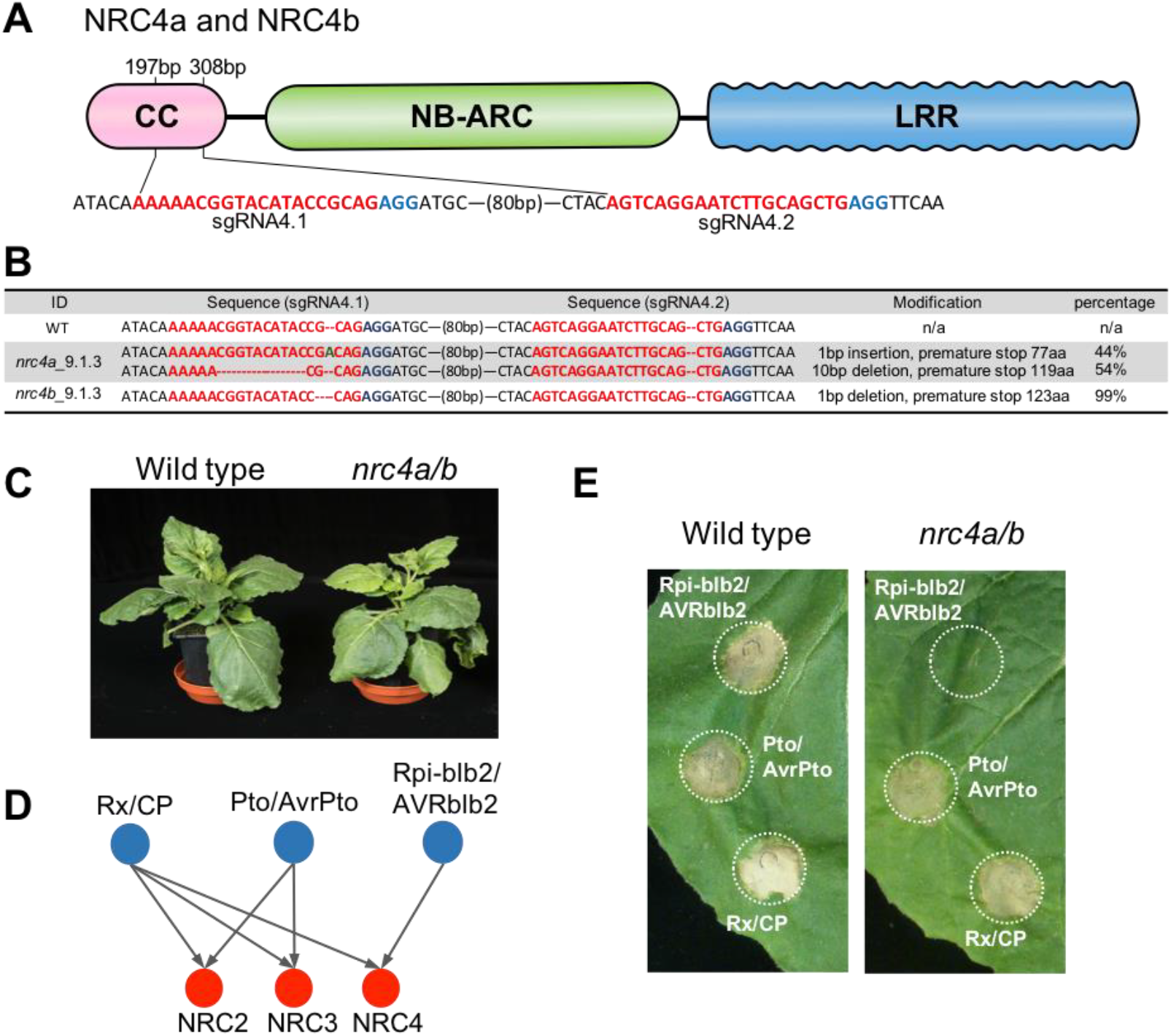
Knocking out of *NRC4a* and *NRC4b* in *Nicotiana benthamiana* impairs Rpi-blb2-mediated HR cell death. (**A**) Schematic representation of sgRNA positions targeting *NRC4a* and *NRC4b*. The PAM motifs are marked in blue, and the sequences of sgRNAs are marked in red. (**B**) Genotyping results of selected T2 *nrc4a/b* plants. Sequences of the two sgRNA positions in *NRC4a* and *NRC4b* were confirmed by amplicon sequencing. “Percentage” represents the proportion of Illumina reads belonging to each sequence category. (**C**) *NRC4* knockout lines did not exhibit any growth defects when compared to wild type plants. Photographs of five weeks old wild type and *nrc4ab* knock out *N. benthamiana* plants. (**D**) Schematic representation showing the genetic dependency of Rpi-blb2, Pto, and Rx on different NRCs according to previous finding with virus-induced gene silencing analysis (Wu et al., 2017). (**E**) Rpi-blb2-mediated HR cell death was compromised in the *NRC4* knockout lines. Rpi-blb2/AVRblb2, Pto/AvrPto and Rx/CP were transiently expressed in leaves of wild type and *nrc4a/b N. benthamiana* according to the method described previously in Wu et al. (2017). The pictures were taken at 7 days after agroinfiltration.

**Figure supplement 2.**
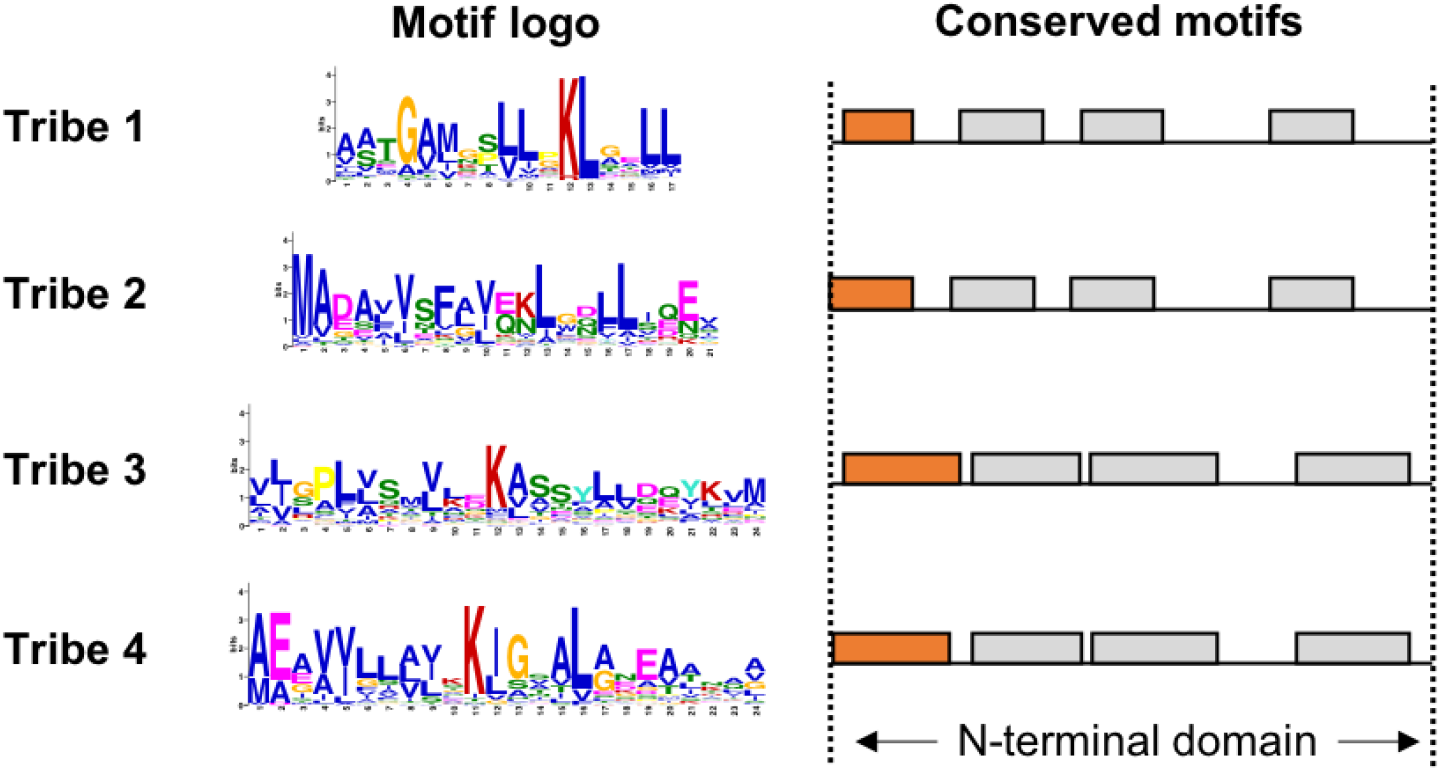
CC-NLRs have conserved protein sequence patterns in the beginning of the N-terminal domains. Consensus sequence patterns in N-terminal domains were identified by MEME from 226 Tribe 1, 102 Tribe 2, 83 Tribe 3 and 59 Tribe 4 members. Motif logos (left) describe the N-terminal consensus patterns from proteins in each tribe, as highlighted in orange (right).

**Figure supplement 3.**
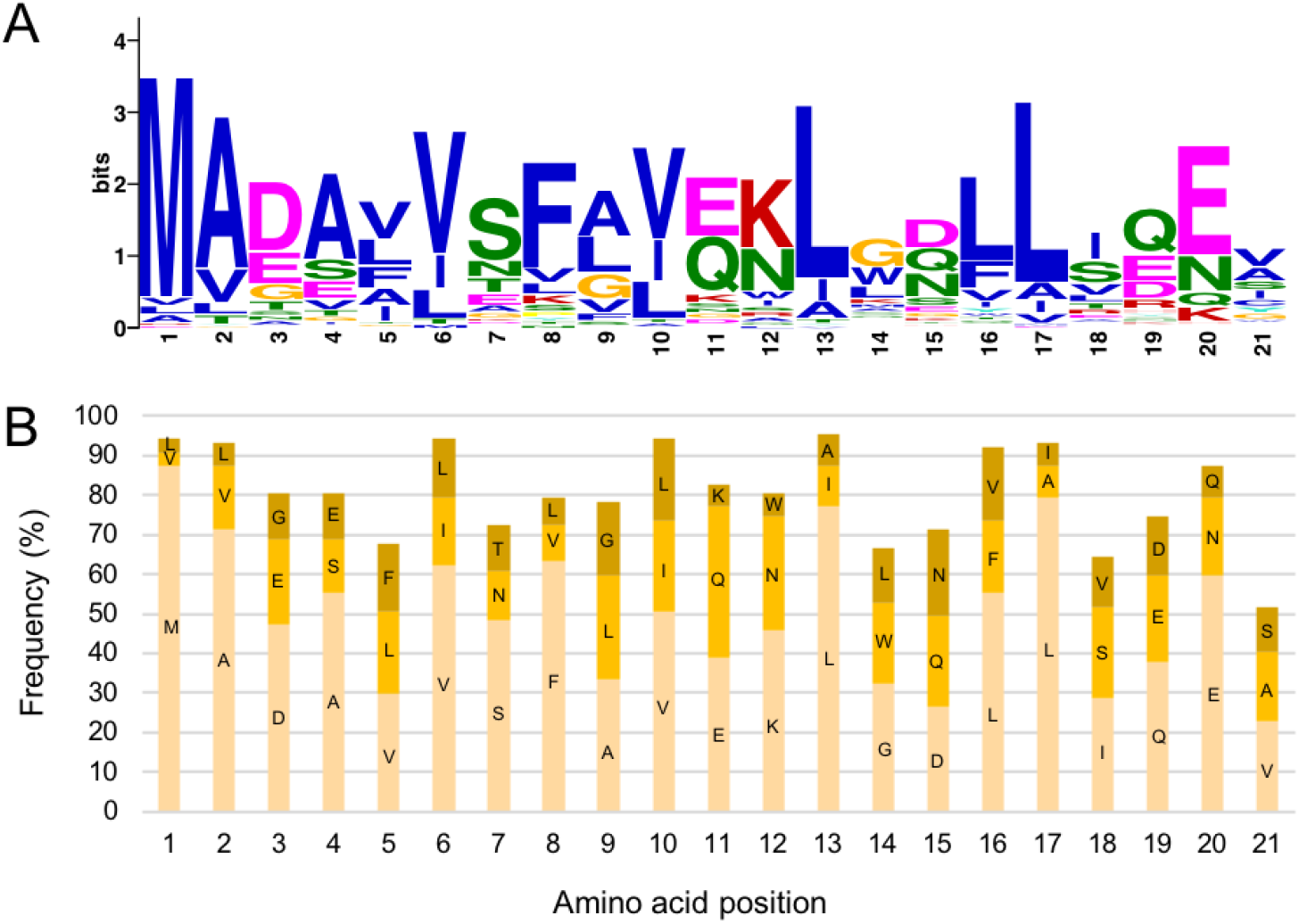
N terminus of NRC4 possesses a consensus pattern coined MADA motif. (**A**) Consensus sequence of the MADA motif. The MADA motif logo was generated by MEME from 87 N-terminal domains of Tribe 2 members. (**B**) Graphical representation of the MADA HMM used to screen MADA-CC-NLR. The three most abundant amino acids at each position in the motif are shown by frequency and labelled with their one-letter code.

**Figure supplement 4.**
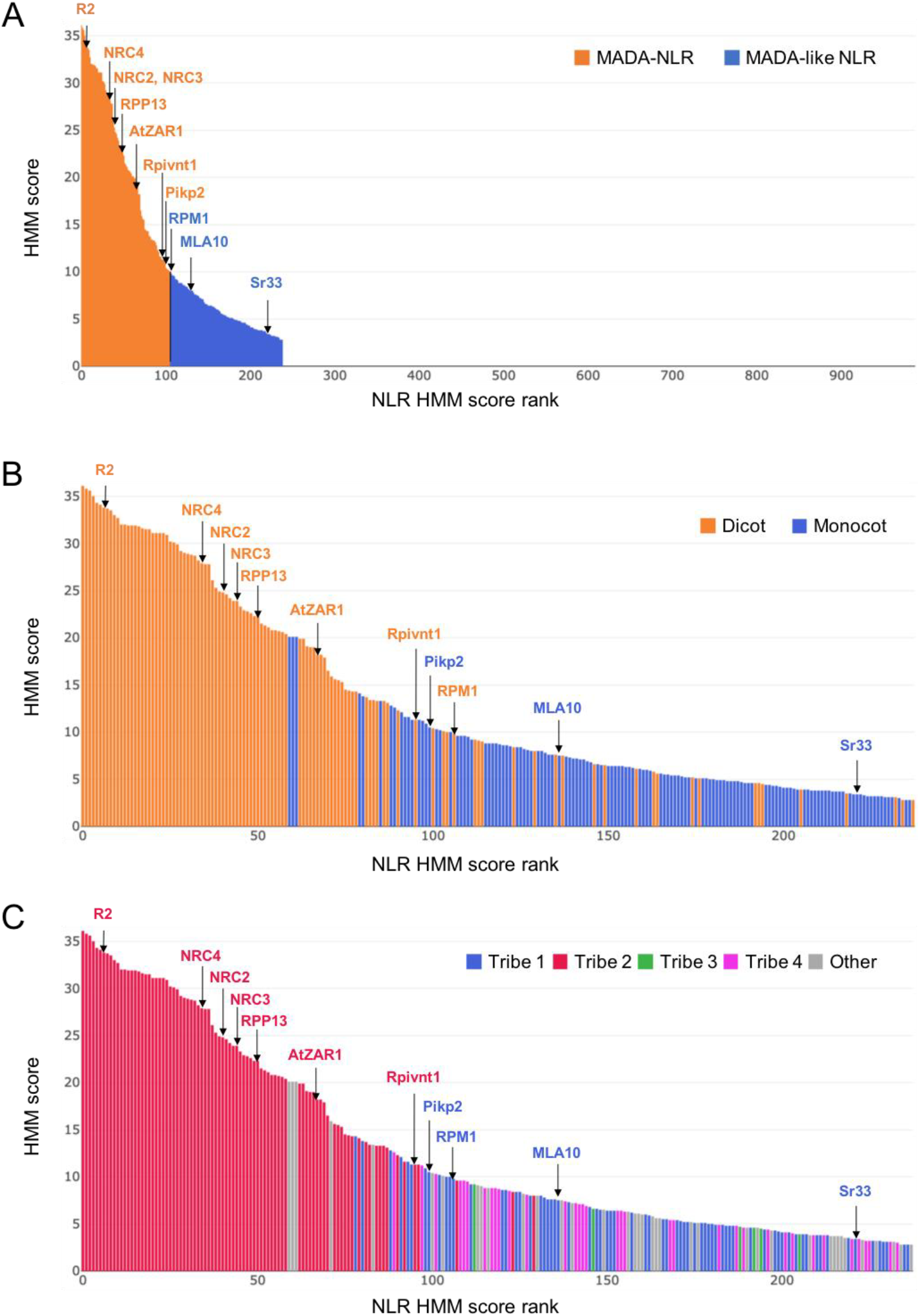
Bar graph of MADA/MADAL-CC-NLRs according to HMM score. (**A**) HMM score bar graph for CC/TIR-NLR database (988 proteins, Supplementary file 2). MADA/MADAL-CC-NLRs from HMMER analysis were shown in orange and blue, respectively. (**B**) HMM score bar graph with plant species information. MADA/MADAL-CC-NLRs from dicot and monocot plant species were presented in orange and blue, respectively. (**C**) HMM score bar graph with N-terminal domain tribe information. MADA/MADAL-CC-NLRs in tribe 1-4 from Tribe-MCL analysis were described in blue, red, green and pink, respectively. MADA/MADAL-CC-NLRs from the other N-terminal domain tribes are shown in grey.

**Figure supplement 5.**
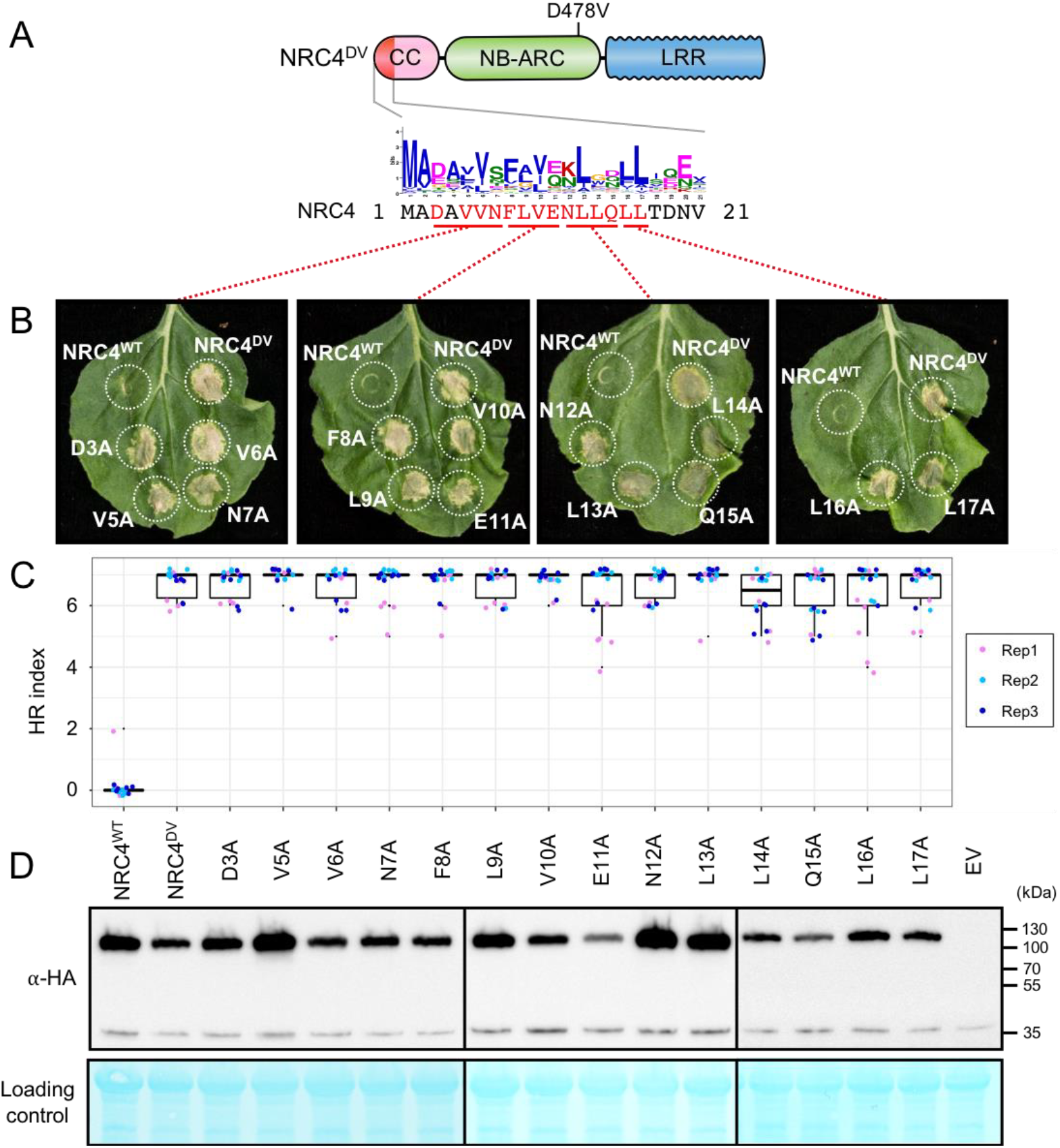
Alanine mutants do not compromise HR cell death triggered by autoactive NRC4. (**A**) Schematic representation of NRC4 and the alanine (A) mutant scan of the MADA motif. Mutated sites are shown as red characters in the NRC4 sequence. (**B**) Cell death observed in *N. benthamiana* after expression of NRC4 mutants. *N. benthamiana* leaf panels expressing NRC4^WT^-6xHA, NRC4^DV^-6xHA and the corresponding A mutants were photographed at 5 days after agroinfiltration. (**C**) Box plots showing cell death intensity scored as an HR index based on three independent experiments. (**D**) *In planta* accumulation of the NRC4 variants. Immunoblot analysis was done as described in Figure 7D.

**Figure supplement 6.**
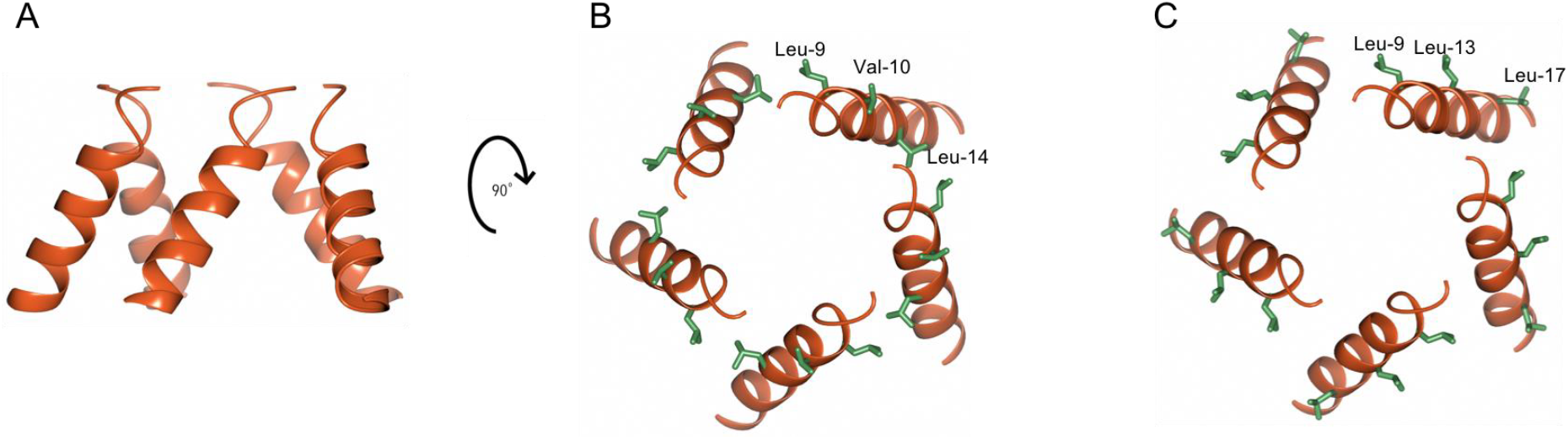
Mapping loss of function mutations on N-terminal α helices of NRC4. (**A**) Cartoon representation of N-terminal α helices of NRC4 resistosome (zoom in grey box of Figure 4B). (**B** and **C**) N-terminal α helices are rotated 90 degrees and mutated amino acids are shown as stick representation and labelled.

**Figure supplement 7.**
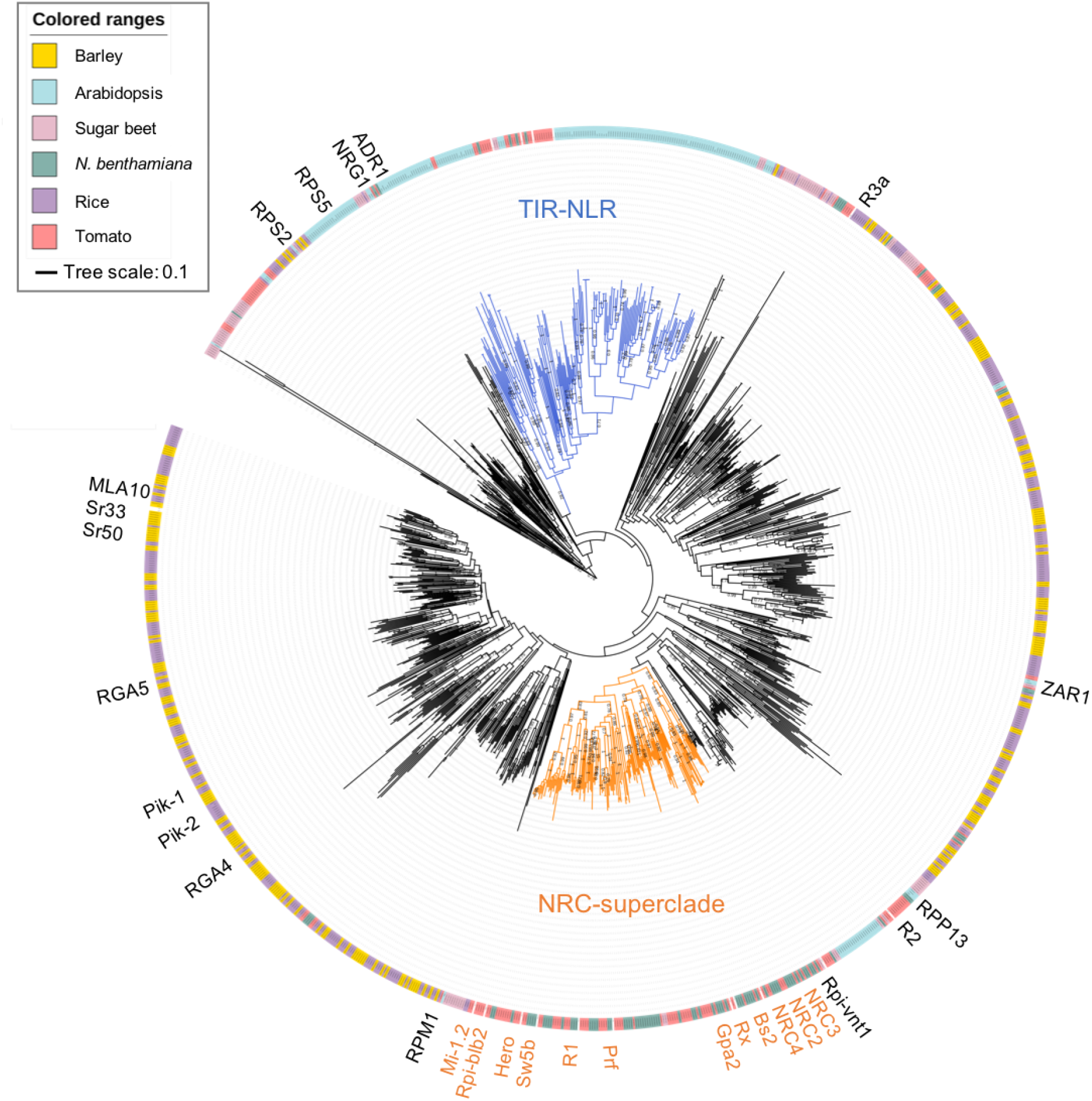
Phylogenetic analysis of NLR proteins from dicot and monocot plant species. NLR proteins were predicted by NLR-parser from *N. benthamiana* (NbS-), tomato (Solyc-), Arabidopsis (AT-), sugar beet (Bv-), rice (Os-) and barley (HORVU-) proteomes, and were used for the MAFFT multiple alignment and phylogenetic analyses. The phylogenetic tree was constructed with the NB-ARC domain sequences in MEGA7 by the neighbour-joining method. Each leaf was labelled with different color ranges indicating plant species. Well-supported TIR-NLR clade and NRC-superclade members were marked in blue and orange, respectively. The bootstrap supports (> 0.7) are indicated as texts. The scale bar indicates the evolutionary distance in amino acid substitution per site.

**Figure supplement 8.**
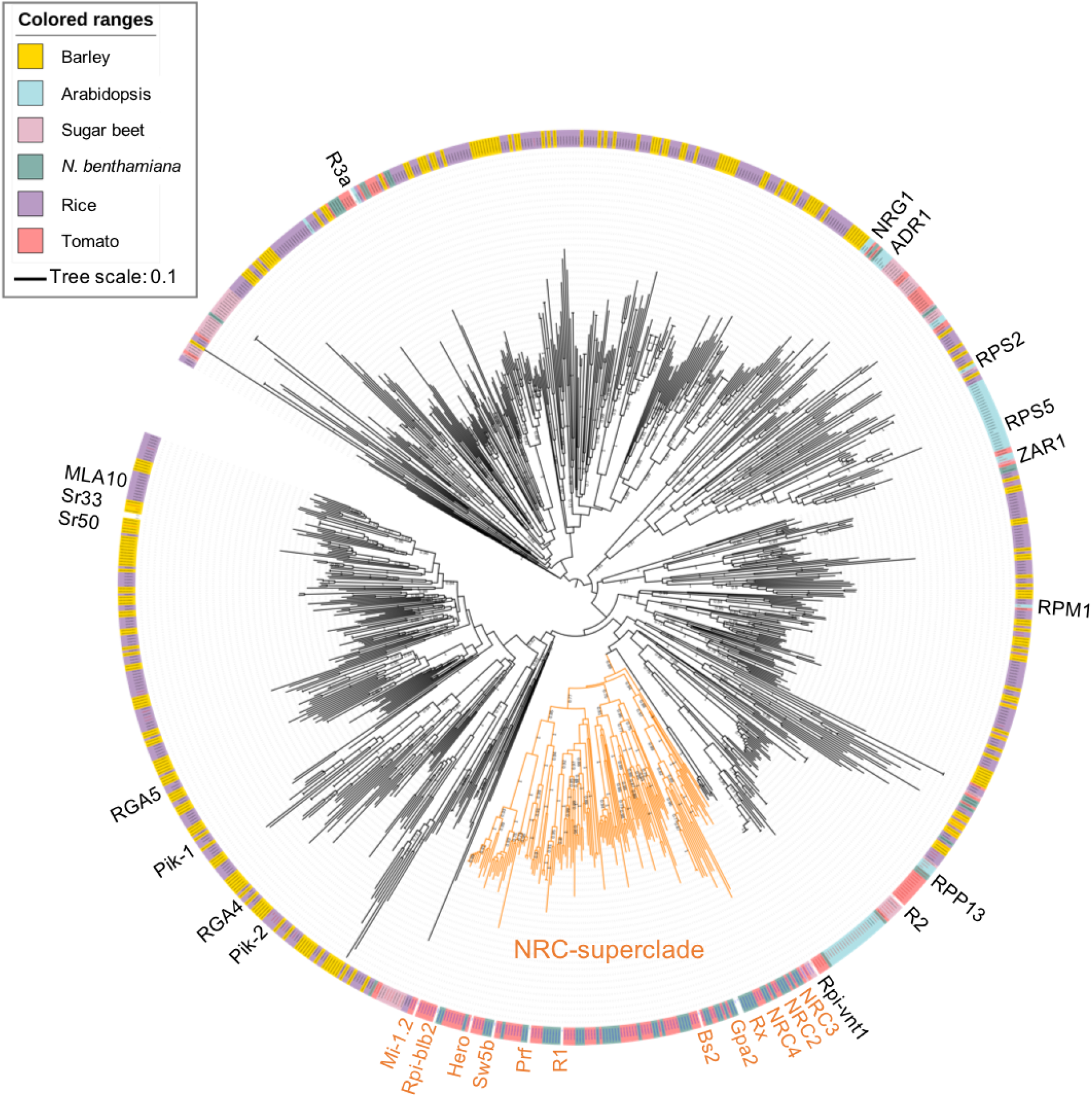
Phylogenetic analysis of CC-NLR proteins from dicot and monocot plant species. The phylogenetic tree was constructed with the NB-ARC domain sequences of CC-NLRs as described in Figure supplement 7. Each leaf was labelled with different color ranges indicating plant species. Well-supported NRC-superclade members were marked in orange. The bootstrap supports (> 0.7) are indicated as texts. The scale bar indicates the evolutionary distance in amino acid substitution per site.

## Supplementary files and tables

**Supplementary file 1. Sequences of NRC4 truncation library.** The Mu-STOP transposon insertion sites were confirmed by PCR amplicon sequencing with Mu-STOP seq Rv primer. The 65 truncate sequences of NRC4 are listed in this file.

**Supplementary file 2. Amino acid sequences of full-length NLRs in the CC-NLR database.** 988 NLR sequences used for HMMER analysis are listed.

**Supplementary file 3. Amino acid sequences of N-terminal domains in the CC-NLR database**. N-terminal domain sequences of 988 proteins used for Tribe-MCL analysis are listed.

**Supplementary file 4. Amino acid sequences for CC/TIR-NLR phylogenetic tree.** NB-ARC domain sequences used for phylogenetic analysis are shown with the IDs, *N. benthamiana* (NbS-), tomato (Solyc-), Arabidopsis (AT-), sugar beet (Bv-), rice (Os-) and barley (HORVU-).

**Supplementary file 5. Amino acid sequences for CC-NLR phylogenetic tree.** NB-ARC domain sequences used for phylogenetic analysis are shown with the IDs, *N. benthamiana* (NbS-), tomato (Solyc-), Arabidopsis (AT-), sugar beet (Bv-), rice (Os-) and barley (HORVU-).

**Supplementary file 6. The MADA-HMM for HMMER analysis.** This MADA-HMM was used for searching MADA-CC-NLRs from CC-NLR database (Supplementary file 2).

**Supplementary Table 1. N-terminal domain tribes of CC-NLRs.** Results of the Tribe-MCL analysis are included in this file.

**Supplementary Table 2. Amino acid sequences of the MADA motif.** The sequences were extracted from MEME output against N-terminal domain tribe 2 and were used to build the MADA motif HMM.

**Supplementary Table 3. Output of the HMMER search using the MADA motif HMM against tomato and Arabidopsis proteomes.** HMM scores are listed with the IDs, tomato (Solyc-) and Arabidopsis (AT-), and annotation information.

**Supplementary Table 4. Output of the HMMER search using the MADA motif HMM against the CC-NLR database.** HMM scores of the predicted MADA motifs are listed by IDs, *N. benthamiana* (NbS-), tomato (Solyc-), Arabidopsis (AT-), sugar beet (Bv-), rice (Os-) and barley (HORVU-), with Tribe-MCL result, the start (‘MADA_strat’) and end (‘MADA_end’) positions of the MADA motifs in the CC-NLRs.

**Supplementary Table 5. HMM scores of NRC-superclade proteins.** HMM scores are listed by IDs, *N. benthamiana* (NbS-), tomato (Solyc-) and sugar beet (Bv-) with Tribe-MCL result, the start (‘MADA_strat’) position of the MADA motifs and NRC clade information (‘NRC-H’ and ‘NRC-S’).

**Supplementary Table 6. List of the predicted Arabidopsis MADA-CC-NLRs.** The IDs are listed with the HMM score.

**Supplementary Table 7.** Primers used for generating NRC4 variants by Golden Gate cloning.

