## Supplementary files and tables for "An N-terminal motif in NLR immune receptors is functionally conserved across distantly related plant species": Supplementary table 7.pptx

### Slide 1
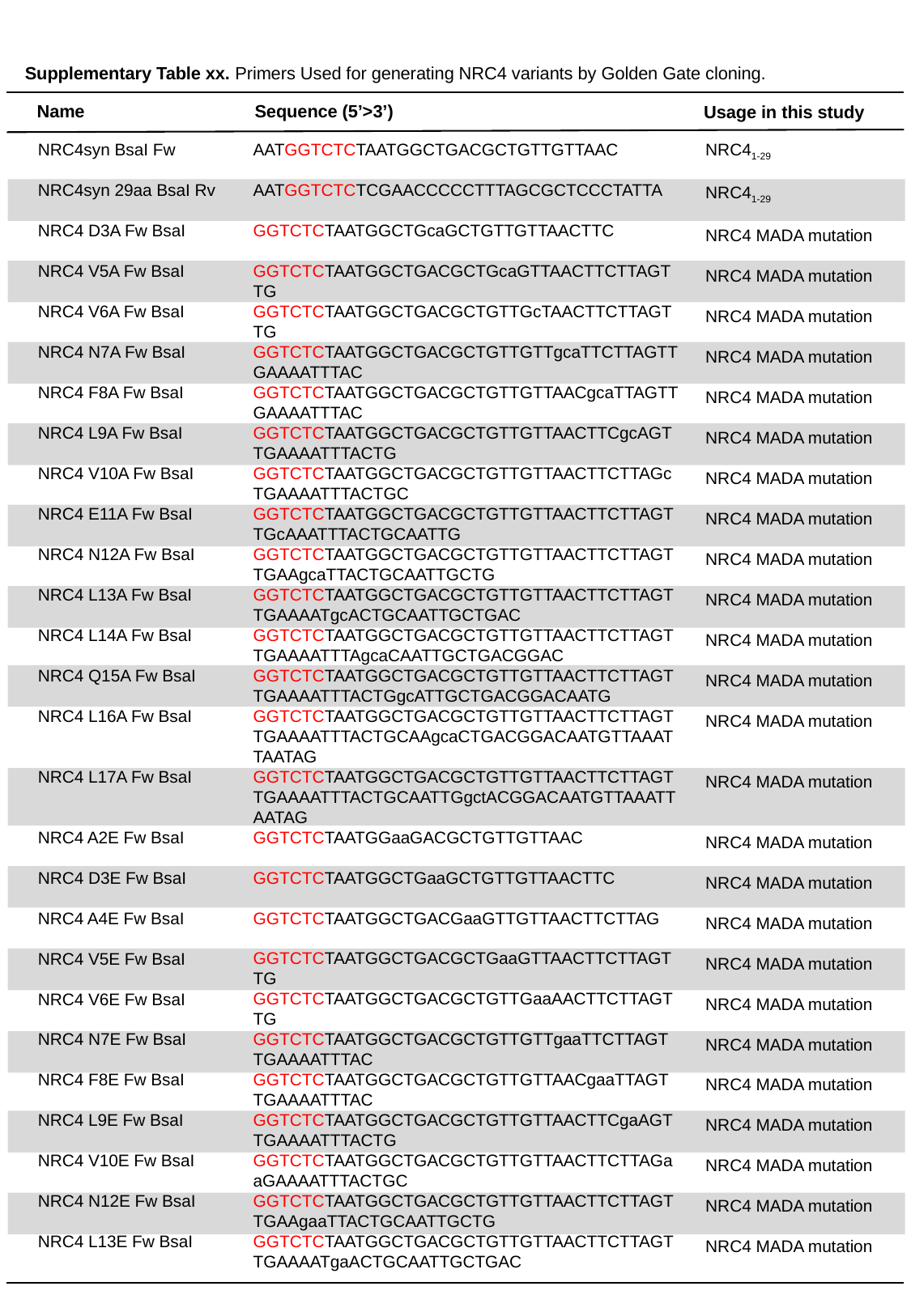

Supplementary Table xx. Primers Used for generating NRC4 variants by Golden Gate cloning.
Name
Sequence (5’>3’)
Usage in this study
NRC4_1-29_
NRC4_1-29_
NRC4 MADA mutation
NRC4 MADA mutation
NRC4 MADA mutation
NRC4 MADA mutation
NRC4 MADA mutation
NRC4 MADA mutation
NRC4 MADA mutation
NRC4 MADA mutation
NRC4 MADA mutation
NRC4 MADA mutation
NRC4 MADA mutation
NRC4 MADA mutation
NRC4 MADA mutation
NRC4 MADA mutation
NRC4 MADA mutation
NRC4 MADA mutation
NRC4 MADA mutation
NRC4 MADA mutation
NRC4 MADA mutation
NRC4 MADA mutation
NRC4 MADA mutation
NRC4 MADA mutation
NRC4 MADA mutation
NRC4 MADA mutation
NRC4 MADA mutation
NRC4syn BsaI Fw
NRC4syn 29aa BsaI Rv
NRC4 D3A Fw BsaI
NRC4 V5A Fw BsaI
NRC4 V6A Fw BsaI
NRC4 N7A Fw BsaI
NRC4 F8A Fw BsaI
NRC4 L9A Fw BsaI
NRC4 V10A Fw BsaI
NRC4 E11A Fw BsaI
NRC4 N12A Fw BsaI
NRC4 L13A Fw BsaI
NRC4 L14A Fw BsaI
NRC4 Q15A Fw BsaI
NRC4 L16A Fw BsaI
NRC4 L17A Fw BsaI
NRC4 A2E Fw BsaI
NRC4 D3E Fw BsaI
NRC4 A4E Fw BsaI
NRC4 V5E Fw BsaI
NRC4 V6E Fw BsaI
NRC4 N7E Fw BsaI
NRC4 F8E Fw BsaI
NRC4 L9E Fw BsaI
NRC4 V10E Fw BsaI
NRC4 N12E Fw BsaI
NRC4 L13E Fw BsaI
AATGGTCTCTAATGGCTGACGCTGTTGTTAAC
AATGGTCTCTCGAACCCCCTTTAGCGCTCCCTATTA
GGTCTCTAATGGCTGcaGCTGTTGTTAACTTC
GGTCTCTAATGGCTGACGCTGcaGTTAACTTCTTAGTTG
GGTCTCTAATGGCTGACGCTGTTGcTAACTTCTTAGTTG
GGTCTCTAATGGCTGACGCTGTTGTTgcaTTCTTAGTTGAAAATTTAC
GGTCTCTAATGGCTGACGCTGTTGTTAACgcaTTAGTTGAAAATTTAC
GGTCTCTAATGGCTGACGCTGTTGTTAACTTCgcAGTTGAAAATTTACTG
GGTCTCTAATGGCTGACGCTGTTGTTAACTTCTTAGcTGAAAATTTACTGC
GGTCTCTAATGGCTGACGCTGTTGTTAACTTCTTAGTTGcAAATTTACTGCAATTG
GGTCTCTAATGGCTGACGCTGTTGTTAACTTCTTAGTTGAAgcaTTACTGCAATTGCTG
GGTCTCTAATGGCTGACGCTGTTGTTAACTTCTTAGTTGAAAATgcACTGCAATTGCTGAC
GGTCTCTAATGGCTGACGCTGTTGTTAACTTCTTAGTTGAAAATTTAgcaCAATTGCTGACGGAC
GGTCTCTAATGGCTGACGCTGTTGTTAACTTCTTAGTTGAAAATTTACTGgcATTGCTGACGGACAATG
GGTCTCTAATGGCTGACGCTGTTGTTAACTTCTTAGTTGAAAATTTACTGCAAgcaCTGACGGACAATGTTAAATTAATAG
GGTCTCTAATGGCTGACGCTGTTGTTAACTTCTTAGTTGAAAATTTACTGCAATTGgctACGGACAATGTTAAATTAATAG
GGTCTCTAATGGaaGACGCTGTTGTTAAC
GGTCTCTAATGGCTGaaGCTGTTGTTAACTTC
GGTCTCTAATGGCTGACGaaGTTGTTAACTTCTTAG
GGTCTCTAATGGCTGACGCTGaaGTTAACTTCTTAGTTG
GGTCTCTAATGGCTGACGCTGTTGaaAACTTCTTAGTTG
GGTCTCTAATGGCTGACGCTGTTGTTgaaTTCTTAGTTGAAAATTTAC
GGTCTCTAATGGCTGACGCTGTTGTTAACgaaTTAGTTGAAAATTTAC
GGTCTCTAATGGCTGACGCTGTTGTTAACTTCgaAGTTGAAAATTTACTG
GGTCTCTAATGGCTGACGCTGTTGTTAACTTCTTAGaaGAAAATTTACTGC
GGTCTCTAATGGCTGACGCTGTTGTTAACTTCTTAGTTGAAgaaTTACTGCAATTGCTG
GGTCTCTAATGGCTGACGCTGTTGTTAACTTCTTAGTTGAAAATgaACTGCAATTGCTGAC

### Slide 2
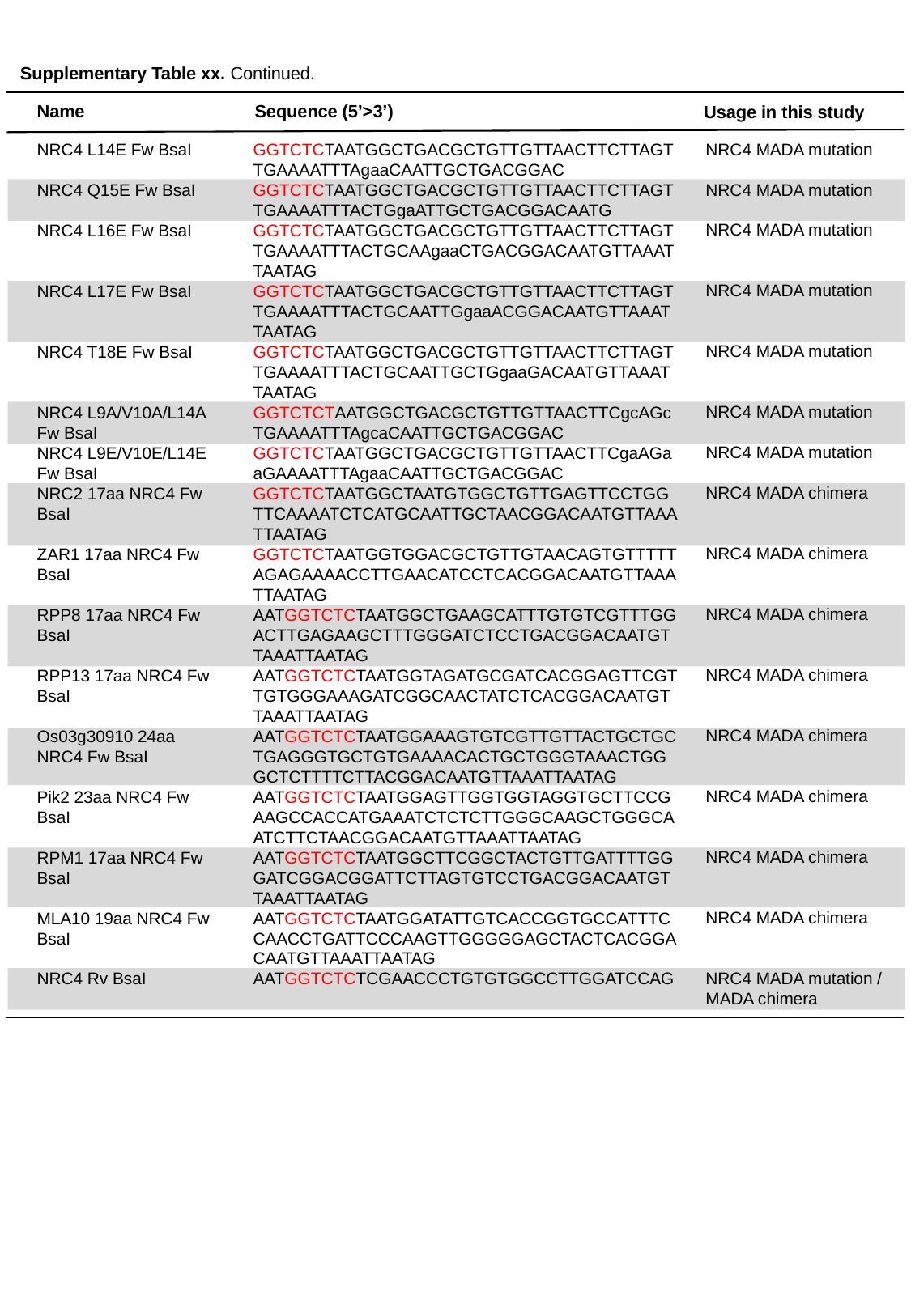

Supplementary Table xx. Continued.
Name
Sequence (5’>3’)
Usage in this study
NRC4 MADA mutation
NRC4 MADA mutation
NRC4 MADA mutation
NRC4 MADA mutation
NRC4 MADA mutation
NRC4 MADA mutation
NRC4 MADA mutation
NRC4 MADA chimera
NRC4 MADA chimera
NRC4 MADA chimera
NRC4 MADA chimera
NRC4 MADA chimera
NRC4 MADA chimera
NRC4 MADA chimera
NRC4 MADA chimera
NRC4 MADA mutation /
MADA chimera
NRC4 L14E Fw BsaI
NRC4 Q15E Fw BsaI
NRC4 L16E Fw BsaI
NRC4 L17E Fw BsaI
NRC4 T18E Fw BsaI
NRC4 L9A/V10A/L14A Fw BsaI
NRC4 L9E/V10E/L14E Fw BsaI
NRC2 17aa NRC4 Fw BsaI
ZAR1 17aa NRC4 Fw BsaI
RPP8 17aa NRC4 Fw BsaI
RPP13 17aa NRC4 Fw BsaI
Os03g30910 24aa NRC4 Fw BsaI
Pik2 23aa NRC4 Fw BsaI
RPM1 17aa NRC4 Fw BsaI
MLA10 19aa NRC4 Fw BsaI
NRC4 Rv BsaI
GGTCTCTAATGGCTGACGCTGTTGTTAACTTCTTAGTTGAAAATTTAgaaCAATTGCTGACGGAC
GGTCTCTAATGGCTGACGCTGTTGTTAACTTCTTAGTTGAAAATTTACTGgaATTGCTGACGGACAATG
GGTCTCTAATGGCTGACGCTGTTGTTAACTTCTTAGTTGAAAATTTACTGCAAgaaCTGACGGACAATGTTAAATTAATAG
GGTCTCTAATGGCTGACGCTGTTGTTAACTTCTTAGTTGAAAATTTACTGCAATTGgaaACGGACAATGTTAAATTAATAG
GGTCTCTAATGGCTGACGCTGTTGTTAACTTCTTAGTTGAAAATTTACTGCAATTGCTGgaaGACAATGTTAAATTAATAG
GGTCTCTAATGGCTGACGCTGTTGTTAACTTCgcAGcTGAAAATTTAgcaCAATTGCTGACGGAC
GGTCTCTAATGGCTGACGCTGTTGTTAACTTCgaAGaaGAAAATTTAgaaCAATTGCTGACGGAC
GGTCTCTAATGGCTAATGTGGCTGTTGAGTTCCTGGTTCAAAATCTCATGCAATTGCTAACGGACAATGTTAAATTAATAG
GGTCTCTAATGGTGGACGCTGTTGTAACAGTGTTTTTAGAGAAAACCTTGAACATCCTCACGGACAATGTTAAATTAATAG
AATGGTCTCTAATGGCTGAAGCATTTGTGTCGTTTGGACTTGAGAAGCTTTGGGATCTCCTGACGGACAATGTTAAATTAATAG
AATGGTCTCTAATGGTAGATGCGATCACGGAGTTCGTTGTGGGAAAGATCGGCAACTATCTCACGGACAATGTTAAATTAATAG
AATGGTCTCTAATGGAAAGTGTCGTTGTTACTGCTGCTGAGGGTGCTGTGAAAACACTGCTGGGTAAACTGGGCTCTTTTCTTACGGACAATGTTAAATTAATAG
AATGGTCTCTAATGGAGTTGGTGGTAGGTGCTTCCGAAGCCACCATGAAATCTCTCTTGGGCAAGCTGGGCAATCTTCTAACGGACAATGTTAAATTAATAG
AATGGTCTCTAATGGCTTCGGCTACTGTTGATTTTGGGATCGGACGGATTCTTAGTGTCCTGACGGACAATGTTAAATTAATAG
AATGGTCTCTAATGGATATTGTCACCGGTGCCATTTCCAACCTGATTCCCAAGTTGGGGGAGCTACTCACGGACAATGTTAAATTAATAG
AATGGTCTCTCGAACCCTGTGTGGCCTTGGATCCAG
